# Extracellular disintegration of viral proteins as an innovative strategy for developing broad-spectrum antivirals against coronavirus

**DOI:** 10.1101/2022.11.24.517008

**Authors:** Ke Sun, Zhe Ding, Xiaoying Jia, Haonan Cheng, Yingwen Li, Yan Wu, Zhuoyu Li, Xiaohua Huang, Fangxu Pu, Entao Li, Guiyou Wang, Wei Wang, Yun Ding, Gary Wong, Sandra Chiu, Jiaming Lan, Aiguo Hu

**Affiliations:** Shanghai Key Laboratory of Advanced Polymeric Materials, School of Materials Science and Engineering, East China University of Science and Technology, Shanghai 200237, China; The Institut Pasteur of Shanghai, Chinese Academy of Sciences, Shanghai 200031, China; State Key Laboratory of Virology, Center for Biosafety Mega-Science, Wuhan Institute of Virology, Chinese Academy of Sciences, Wuhan 430071, China; University of the Chinese Academy of Sciences, Beijing 100049, China; Division of Life Sciences and Medicine, University of Science and Technology of China, Hefei, Anhui 230027, China; Department of Clinical Laboratory, The First Affiliated Hospital of USTC, Division of Life Sciences and Medicine, University of Science and Technology of China, Hefei, Anhui 230001, China

**Keywords:** Enediynes, Antiviral agents, Broad-spectrum, Coronavirus, SARS-CoV-2

## Abstract

The coronavirus disease 2019 (COVID-19) pandemic caused by severe acute respiratory syndrome coronavirus 2 (SARS-CoV-2) has claimed millions of lives worldwide, not to mention innumerable losses in the global economy and disruptions in social relationships. Unfortunately, state-of-the-art treatments still lag behind the fast emergence of new variants of concern. The key to resolve this issue is to develop broad-spectrum antivirals with innovative antiviral mechanisms in which coronaviruses are deactivated regardless of their variant development. Herein, we report a new antiviral strategy involving extracellular disintegration of viral proteins that are indispensable for viral infection with hyperanion-grafted enediyne molecules. The sulfate groups ensure low cellular permeability and rather low cytotoxicity of the molecules, while the core enediyne generates reactive radical species and causes significant damage to the spike (S) protein of coronavirus. The enediyne compounds exhibit antiviral activity at micromolar to nanomolar concentrations, and the selectivity index of up to 20,000 against four kinds of human coronaviruses, including the SARS-CoV-2 omicron variant, suggesting the high potential of this new strategy in combating the COVID-19 pandemic.

---

The ongoing coronavirus disease 2019 (COVID-19) pandemic has caused serious damage to human health, social relationships, and the global economy. Since its first identification in early 2020(*1, 2*), the culprit severe acute respiratory syndrome coronavirus 2 (SARS-CoV-2) has accumulated thousands of mutations(*3*). The World Health Organization (WHO) has listed a series of variants of concern (VOCs) in the last two years, including the predominant alpha, delta and omicron variants that caused significant infection waves worldwide. The rapid mutation of SARS-CoV-2 and fast emergence of new VOCs constitute a great challenge to the colony immune defense by widespread vaccination and antiviral developments, as most of them are designed in the “one bug-one drug” approach. Developing broad-spectrum antivirals, *ideally with an innovative antiviral mechanism*, is therefore of essential importance to combat the current COVID-19 and future “disease X”(*4*) likely caused by other zoonotic viruses as the confronting of the previously distinct ecological niches becomes more frequent(*5*).

SARS-CoV-2 and six other known human coronaviruses are enveloped viruses. They are composed of lipid bilayers and four structural proteins, namely, spike (S), membrane (M), envelope (E) and nucleocapsid (N) proteins, with which the viral genome (approximately 30 kb) is safely wrapped. The S proteins are responsible for the interaction with the receptors on the cell surface to penetrate the host cells and further hijack the cellular reproduction machinery(*6*). Significant efforts have been devoted to developing antivirals against SARS-CoV-2 by blocking the cell entry process using engineered small molecules, proteins, adaptors, and extracellular vesicles to interrupt the binding of S protein to angiotensin converting enzyme 2 (ACE 2) cell receptor(*3, 7-12*). As the glycocalyx on the cell surface is negatively charged(*13, 14*), the electrostatic interaction between glycans and the positively charged receptor binding domains (RBD) and additional binding sites in the S proteins provides localization of the virus for their cellular binding(*15, 16*). To this end, synthetic glycan analogs have been developed as broad-spectrum antivirals. For example, a variety of materials with multiple sulfate (or sulfonate) groups showed prominent results in treating with SARS-CoV-2(*17-19*). Unfortunately, polyanionic systems are anticoagulant in nature, which may increase the risk of off-target adverse effects. Meanwhile, their inhibitory effect toward virus particles might be lost upon dilution in body fluid, resulting in revival of virus infection.

Enediynes, a family of compounds with a (Z)-hexa-3-en-1,5-diyne core structure, first discovered in the 1960s from the culture filtrates of *Streptomyces*(*20*), exert profound biological activity through a hitherto unseen mode of action. After a special triggering mechanism, the activated enediynes undergo Bergman cyclization or Myers-Saito cyclization to generate highly reactive radical species, which further abstract H-atoms from DNA or proteins and show catastrophic effects on tumor cell proliferation(*21*). While natural enediynes are rather scarce and difficult to synthesize(*22*), we recently uncovered a new mechanism, namely, maleimide-assisted rearrangement and cycloaromatization (MARACA, Fig. 1A), enabling the structurally much simpler synthetic enediyne to generate radical species in a physiological environment and exhibit remarkable antitumor activity(*23*). Herein, we report a new strategy for developing broad-spectrum antivirals by installing multiple anionic groups (sulfate groups in this work) onto enediynes. The anionic groups are beneficial for directing the enediyne “warhead” to the S protein of coronavirus through electrostatic interactions. Meanwhile, polyanionic compounds are cell impermeable(*24*), which ensures rather low cytotoxicity to normally functioning cells and the extracellular disintegration of the viral proteins by the radical species generated through MARACA.

**Figure 1.**
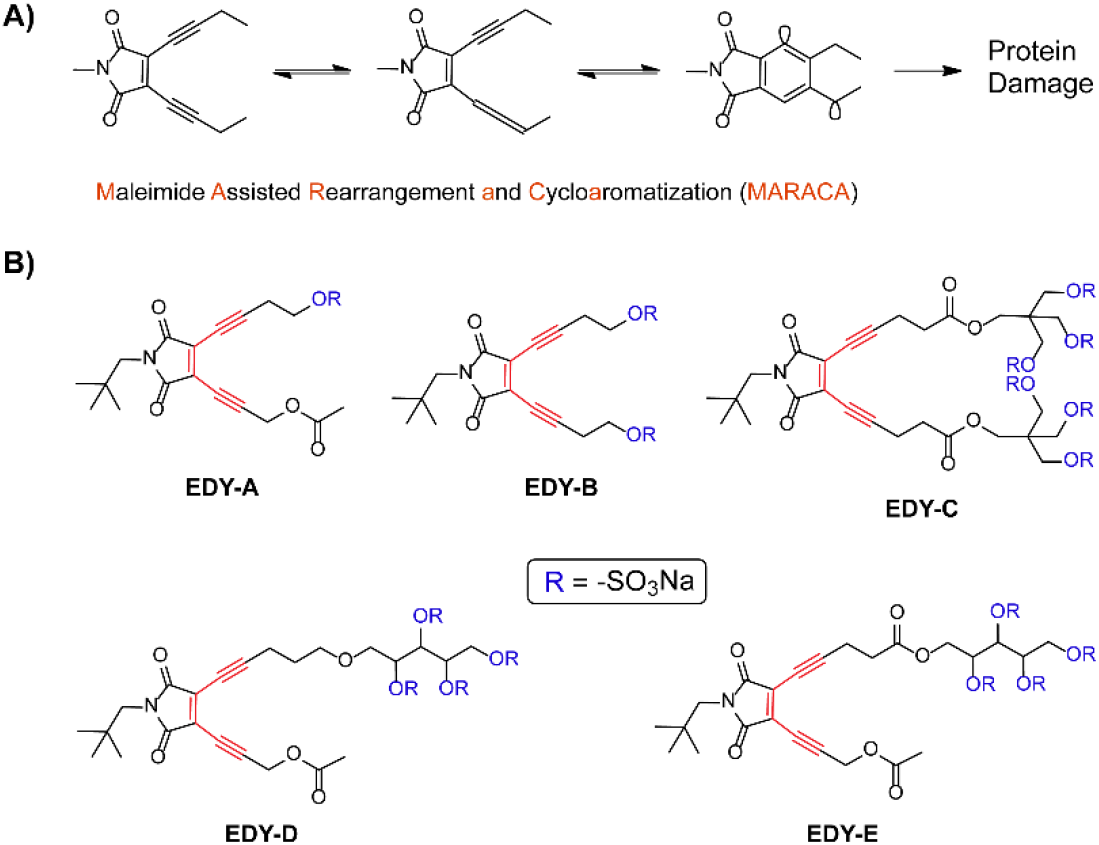
A) Schematic illustration of the radical generation mechanism of maleimide-based enediyne compounds. B) Chemical structures of enediyne antivirals reported in this work.

Figure 1B shows the chemical structures of the sulfated enediynes (**EDY-A** ∼ **EDY-E**), and the detailed synthetic procedures and structural characterizations for these compounds can be found in the supplementary material. Starting from commercially available reagents, these enediynes were synthesized in the longest linear sequence (LLS) of no more than five steps, suggesting their ready scaling-up for future mass production. The first two steps for the preparation of 3,4-diiodomaleimide (**2**) were well documented in our previous work(*25*). The core enediyne structures were constructed in the third step through a Sonogashira coupling reaction between **2** and terminal alkynes, which are either commercially available or synthesized in less than three steps. The enediynes with 1, 2, 4, and 6 hydroxy groups were then hypersulfated in N,N-dimethylformamide (DMF) with excess SO_3_-DMF and neutralized with cold sodium bicarbonate solution followed by removal of inorganic salts and lyophilization to give **EDY-A** ∼ **EDY-E** in high yields. For the synthesis of **EDY-D** and **EDY-E**, an extra step was needed to remove the ketal protection groups before hypersulfation.

The radical-generating property of the enediynes was characterized with electron paramagnetic resonance (EPR) spectroscopy by exploiting a radical trapping strategy. Taking **EDY-A** and **EDY-B** as examples, the appearance of triplet doublet signals in the EPR spectra (Fig. 2A) suggests the generation of radical species from enediynes followed by trapping with N-tert-butyl-α-phenylnitrone (PBN)(*23, 25, 26*). It is noteworthy that under the same conditions, the peaks shown in the EPR spectrum of **EDY-A** are more profound than those for **EDY-B**. We recently confirmed that the introduction of an oxygen atom at the propargyl position of one alkynyl arm of enediyne compounds (such as the one in **EDY-A**) leads to a much faster reaction due to the facilitation of a cascade rearrangement and cycloaromatization with an approximately 6 kcal/mol lower energy barrier(*27*).

**Figure 2.**
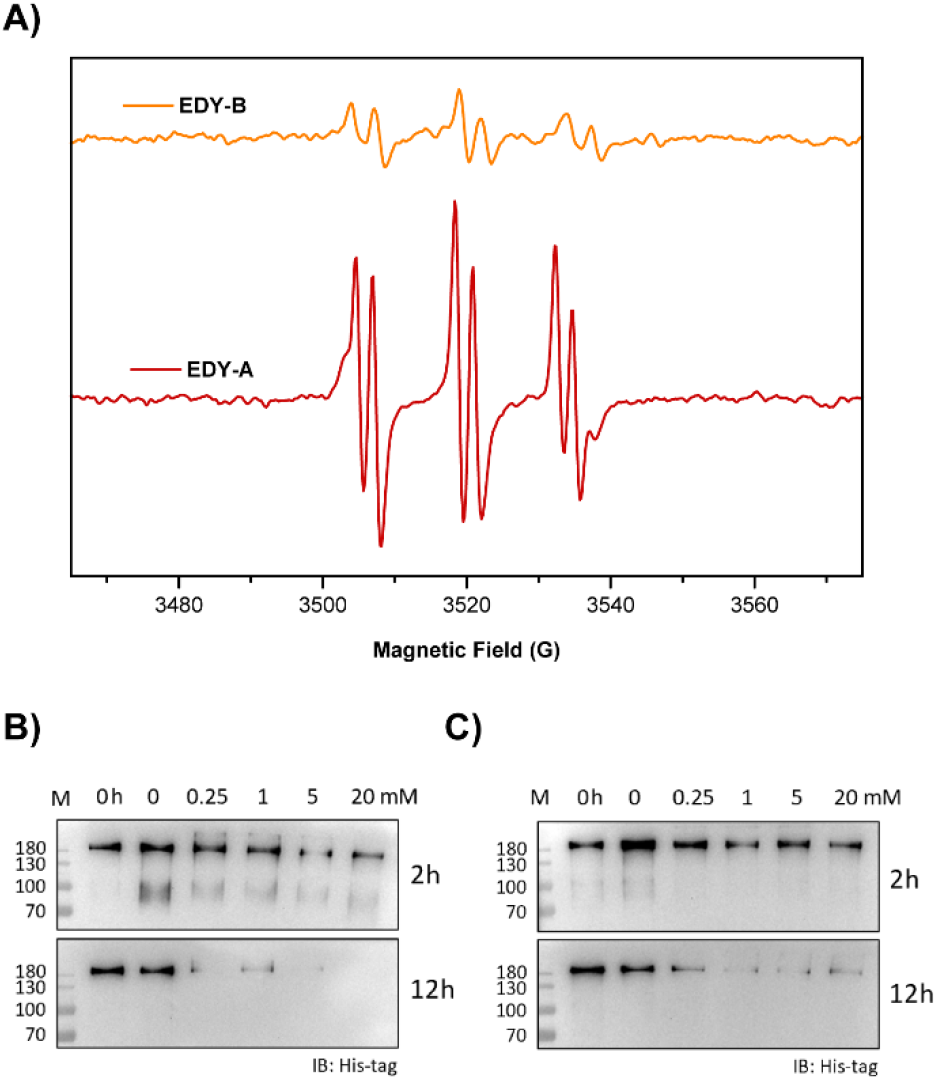
A) EPR spectra of EDY-A and EDY-B after treatment with the spin trapping agent PBN. B) and C) Western blot analysis results of the SARS-CoV-2 S protein after incubation with EDY-A or EDY-B, respectively. The lines marked with “0 h” indicate the standard S protein samples.

The highly reactive radicals generated from enediynes abstract hydrogen atoms from the protein backbone and lead to oxidative disintegration of protein structure and functionality(*28*). To test the ability of enediynes to destroy viral structural proteins, the S protein of SARS-CoV-2 was incubated at 37°C with gradient concentrations of enediynes. Western blot analysis results (Fig. 2B) indicate that the disintegration of S protein is concentration and time dependent. At a concentration of 0.25 mM, **EDY-A** led to significant degradation of the S protein within 2 h of incubation, and further degradation of the S protein was observed at longer incubation time. In comparison, **EDY-B** shows a lower protein disintegration ability at the same time interval, which is in line with its slower radical generation rate, as mentioned above. The protein disintegration results of **EDY-C**∼**EDY-E** exhibited a similar trend and are shown in the supplementary material (Fig. S1). To further confirm the radical nature of the disintegration of viral protein by enediyne, S protein was coincubated with **EDY-D** (5 mM) and serial dilutions of a radical scavenger vitamin C(*29*). At a high concentration of the radical scavenger, the S protein remained intact after incubation at 37°C for 12 h. When the concentration of the radical scavenger was comparable to or lower than that of **EDY-D**, the breakdown or even complete disintegration of the S protein was clearly observed (Fig. S2). The structural integrity of the S protein is essential for viral infection(*30, 31*). Therefore, disintegrating the S protein of coronavirus with enediyne compounds provides a straightforward way to diminish infectivity and eventually deactivate viral particles.

The extremely high cytotoxicity has been the hallmark of enediyne antitumor antibiotics since their early discovery and development. Indeed, two kinds of antibody drug conjugates have been approved for the clinical treatment of leukemia with enediynes as the “warheads”(*32*). The cytotoxicity of enediynes originates from the intercalation of enediyne into the minor groove of DNA, followed by the generation of radical species, abstraction of hydrogen atoms from DNA, and dysfunction of DNA, which leads to cellular apoptosis. On the other hand, to utilize enediynes as the “warheads” for viral protein disintegration, the cellular entrance process of enediynes must be suppressed to make them benign to normally functioning cells. To this end, the cellular uptake of enediynes with or without sulfate group(s) was studied by confocal laser scanning microscopy (CLSM). As shown in Fig. 3, after grafting with one sulfate group, the internalization of enediyne by HeLa cells was significantly inhibited. For **EDY-B** with two sulfate groups, no cellular uptake was observed. For **EDY-C**∼**EDY-E** with more sulfate groups, no cellular internalization of the enediynes was found (Fig. S3), suggesting no harmful effect of these enediynes to cells.

**Figure 3.**
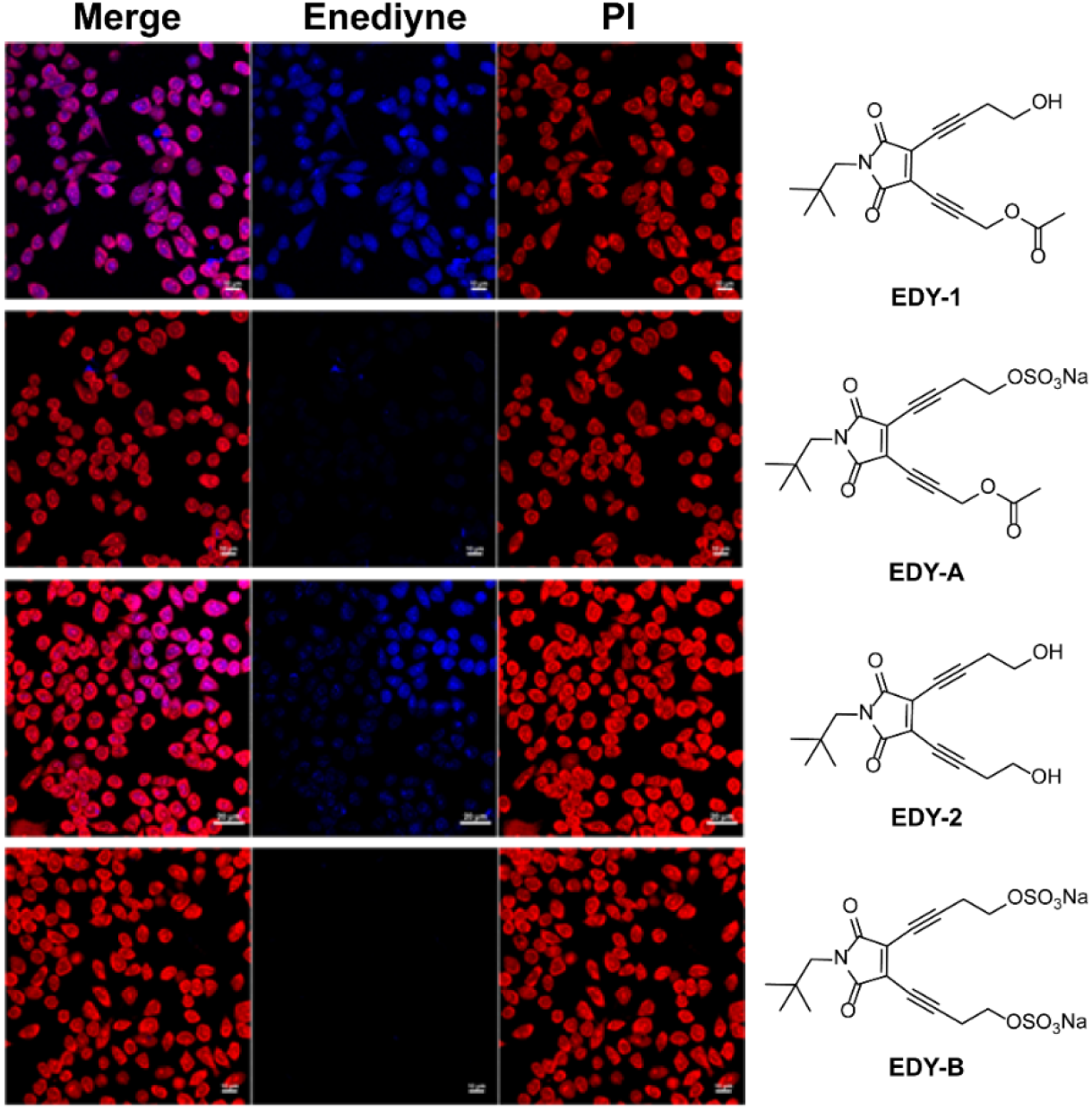
Comparison of the cellular entrance behavior of sulfated enediynes (**EDY-A** and **EDY-B**) with their counterparts (**EDY-1** and **EDY-2**) with confocal laser scanning microscopy. HeLa cells are stained with propidium iodide (PI), showing red fluorescence, while maleimide-based enediynes show intrinsic blue fluorescence(*23*).

With all the information for the extracellular dysfunction of coronavirus in hand, the sulfated enediynes were then evaluated for their ability against human seasonal coronavirus (hCoV-229E, hCoV-NL63 and hCoV-OC43) infections by plaque reduction assay (supplementary materials) in sensitive cell lines (Huh-7, hACE2/Caco2 and RD, respectively). All of the sulfated enediynes showed obvious inhibitory effects with a median effective concentration (EC_50_) value of 3.86 μM∼137.2 μM (Fig. 4). Interestingly, enediynes with oxygen atoms at the propargyl position (**EDY-A, EDY-D** and **EDY-E**) showed high antiviral activities of inhibiting seasonal coronavirus growth with an EC_50_ of approximately 10 μM, while the enediynes (**EDY-B** and **EDY-C**) without this kind of activating functionality(*27*) exhibited an order of magnitude lower antiviral activity, correlating with their relatively slower radical generating rate and weaker viral protein disintegration ability, as discussed above.

**Fig. 4.**
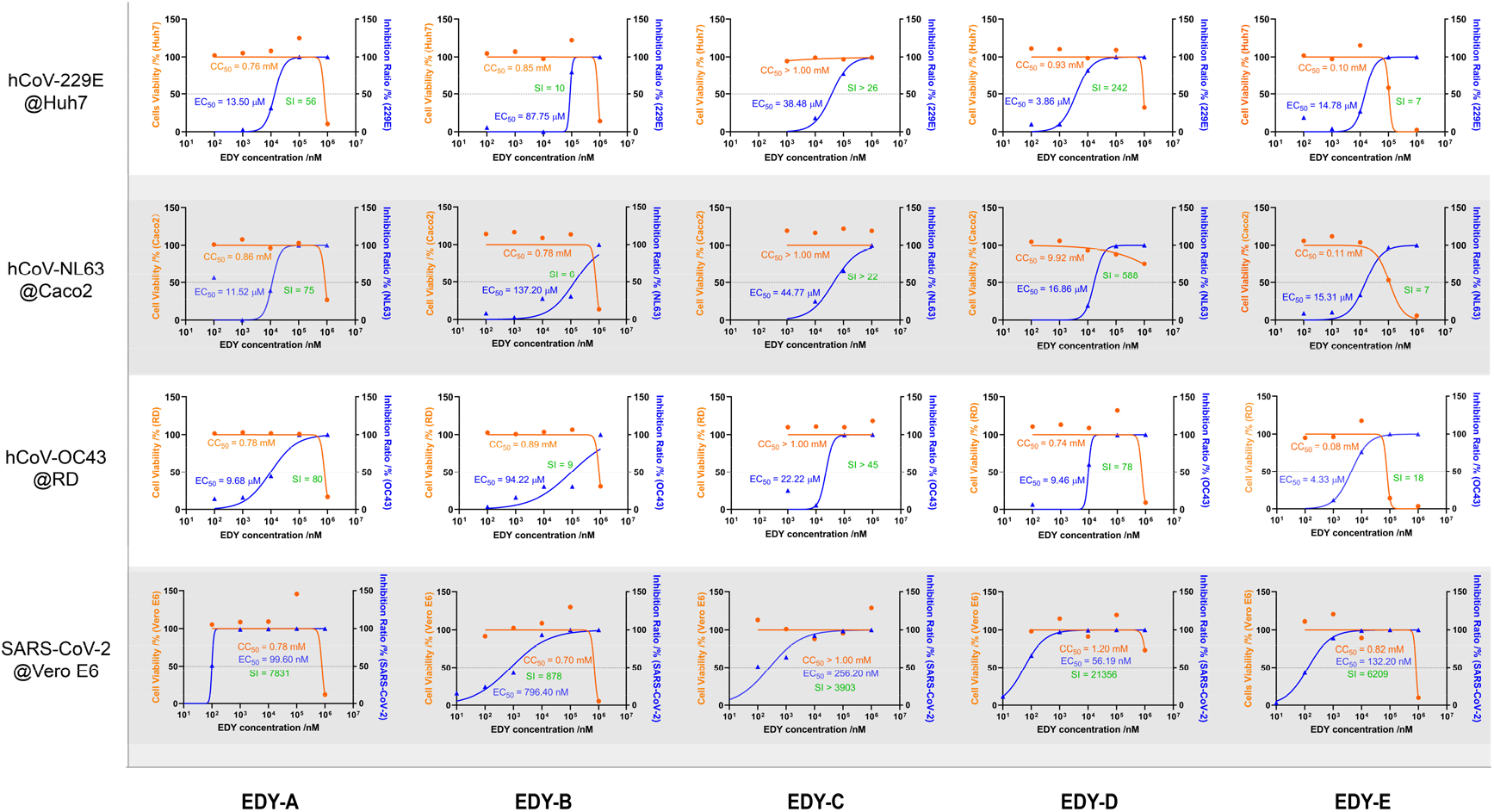
Antiviral activity of enediynes against four kinds of human coronaviruses by plaque reduction assays. Sensitive cell lines were selected for the corresponding coronavirus. The cytotoxicity of these enediynes to different cell lines was measured by CCK-8 assays. The left and right Y-axes of the graphs represent the mean cell variability and inhibition of viral infection in the presence of the enediynes, respectively. A table summarizing all the CC_50_, EC_50_ and SI data can be found in the supplementary materials (Table S1)

The cytotoxicity of enediynes was tested based on a CCK-8 cell counting kit. The Huh-7, hACE2/Caco-2 and RD cells were seeded separately into 96-well microplates (10^4^ cells per well at 100 μL) and cocultured with enediynes at 37 °C (supplementary materials). The good consistency of low cytotoxicity in different cell lines was shown as half cytotoxic concentration (CC_50_) values of approximately 1 mM in **EDY-A, EDY-B** and **EDY-D** (Fig. 4). In addition, **EDY-C** did not exhibit any cytotoxicity even at the highest working concentration (1 mM), which probably benefited from the grafting of six sulfate groups that completely shut down drug entry into the cell to interact with host proteins (and/or DNA). Surprisingly, **EDY-E** showed an order of magnitude higher cytotoxicity even though it is structurally almost the same as **EDY-D**. We speculate that the relatively labile ester linkage in **EDY-E** would lead to the serendipitous release of trace amounts of enediyne core and cause harm to cells due to the high cytotoxicity of the uncharged enediyne(*23*). Altogether, we conclude that the enediyne core is responsible for viral deactivation, while the sulfate groups are indispensable for low cytotoxicity. Both of these parts and the strong linkage between them are essential for antiviral molecular design. For example, **EDY-D** typically showed a high selectivity index (SI, defined as CC_50_/ EC_50_), suggesting a wide application window for potential antiviral treatment.

Unlike human seasonal coronavirus, which typically infects the upper respiratory tract and causes mild symptoms, SARS-CoV-2 can replicate in the lower respiratory tract and is more dangerous, causing much higher morbidity and mortality. The emergence of the VOC omicron and its subvariants with extremely high transmissibility and immune evasion constitute a great challenge to all the current countermeasures(*33-35*). To this end, we tested the performance of **EDY-A**∼**EDY-E** for SARS-CoV-2 omicron treatment. The cytotoxicity experiments of these enediynes to Vero E6 cells were performed regularly, while the antiviral experiments involving SARS-CoV-2 omicron were executed in a biosafety level 3 (BSL-3) laboratory. Encouragingly, all the enediynes showed highly promising antiviral results (Fig. 4). The enediynes with a higher tendency to undergo MARACA and generate radical species typically showed higher antiviral activity. The enediyne with the best performance is **EDY-D**, exhibiting an EC_50_ of 56.19 nM (∼50 ng/mL) and an SI value of over 20,000. The higher antiviral activity of these enediynes toward SARS-CoV-2 omicron than that for the human seasonal viruses is probably due to the enhanced electrostatic interaction between these (poly)sulfated enediynes with the omicron S protein. The omicron S protein shows a positively charged electrostatic surface mainly acquired through N440K, T478K, Q493R, Q498R and Y505H mutations(*34*). While the mutation of the S protein indeed promotes ACE2 recognition, facilitates the infection and transmission of SARS-CoV-2 omicron and is probably responsible for immune evasion of the majority of neutralizing antibodies, when met with these polysulfated enediynes, it goes to the dead end.

In conclusion, we developed an innovative antiviral strategy by employing hyperanion-grafted enediynes to deactivate coronavirus. The core enediyne generates reactive free radical species and causes significant damage to viral proteins and abolishes their function, which is similar to many free radical-generating sanitizing agents. Meanwhile, the anionic groups endow the molecules with rather low cellular permeability and low cytotoxicity and probably guide the enediyne moieties to the RBD region of the S protein, where the peptide chain is positively charged. Overall, this extracellular ultrasanitizing treatment allows disinfection of a broad spectrum of human coronaviruses, including SARS-CoV-2 omicron variants, down to nanomolar concentrations with a high selectivity index for safe use. As the disintegration of viral protein by enediyne is insensitive to the epitopes or subtle structural change of viral proteins, this strategy would also inspire the development of antivirals against other kinds of viruses to fill the huge demand-supply gap(*36*) and might serve as strategic stockpile for combating future “disease X”.

## Acknowledgments

We thank Drs. Junyou Wang and Runhui Liu for the helpful discussions and assistance in the synthesis and biological tests.

## Author contributions

Conceptualization: A.H. and J.L. Synthesis and Characterization: K.S., H.C., Z.L., X.H., and F.P. Cellular Tests: Z.D. and K.S. Antiviral Experiments: Z.D., X.J., Y.W., and E.L. Supervision: A.H., J.L., S.C., and W.W. Funding acquisition: A.H., J.L., S.C., and G.W. Writing-original draft: A.H., J.L., and K.S. Writing-review and editing: Y.D., J.L., A.H., S.C. and G.W.

## Data and materials availability

All data are available in the manuscript or the supplementary materials.

