## supplemental materials for "Extracellular disintegration of viral proteins as an innovative strategy for developing broad-spectrum antivirals against coronavirus"

1. **Materials and Methods for the Synthesis and Analysis of Enediyne Antivirals**
   1.1 **Materials**

Toluene and tetrahydrofuran (THF) were dried using calcium hydride and distilled before use. Anhydrous N,N-dimethylformamide (DMF) with molecular sieves (Water ≤ 30 ppm) was provided by Energy Chemical. Dialysis bag MD31 (MW100-500) were obtained from Beijing Solarbio Science & Technology Co., Ltd. Other reagents were purchased at commercial grade and used without further purification. Sonogashira reactions were performed with dry Schlenk techniques under an atmosphere of nitrogen.

1.2 **Methods**

^1^H NMR (600 MHz) and ^13^C NMR (151 MHz) spectra were recorded on an Ultra Shield 600 spectrometer (BRUKER BIOSPIN AG, AVANCE III 600) and referenced to Me_4_Si. Chemical shifts (δ) are reported in ppm via the deuterated solvent peaks (chloroform, methanol and water) as the standard. Multiplicities were abbreviated as s for singlet, d for doublet, t for triplet and m for multiplet. Mass spectra was recorded on a Micromass LCTTM mass spectrometer using the ESI method.

1.3 **Synthesis**

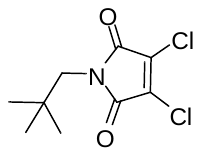

3,4-dichloro-1-(2,2-dimethyl-propyl)-pyrrole-2,5-dione (**1**)

This compound was synthesized following the procedure described in our previous work(*1*). Briefly, dichloromaleic anhydride (7.95 g, 47.9 mmol) was dissolved in acetic acid (50 mL) with slow addition of neopentylamine (3.80 g, 43.55 mmol) at 0 °C, and then the solution was heated at 120 °C for 18 h. After removal of the solvent, the crude residue was separated by column chromatography on silica gel (ethyl acetate / hexane = 1:19) to give the product (8.5 g, 82%).

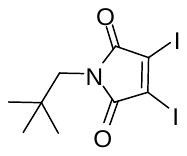

3,4-diiodo-1-(2,2-dimethyl-propyl)-pyrrole-2,5-dione (**2**)

This compound was synthesized following the procedure described in our previous work (*1*). Briefly, a mixture of sodium iodide (40.9 g, 272.8 mmol) and **1** (16.1 g, 68.2 mmol) in acetonitrile (140 mL) was heated at 85 °C for 24 h. After removal of the solvent, the residue was purified by column chromatography on silica gel (dichloromethane, DCM) to give the product (26.7 g, 93%). ^1^H NMR (CDCl_3_) δ 3.43 (s, 2H), 0.92 (s, 9H).

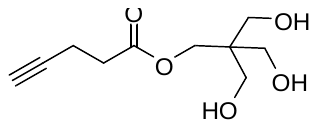

3-hydroxy-2,2-bis(hydroxymethyl) propyl pent-4-ynoate (**3**)

This compound was synthesized according to a reported procedure with slight modification(*2*). Briefly, 4-pentynoic acid (2.16 g, 22 mmol), pentaerythritol (6.0 g, 44 mmol), and 4-dimethylaminopyridine (537 mg, 4.4 mmol) were fully dissolved in anhydrous DMF (40 mL). The flask was cooled to 0 °C and stirred for 10 min, followed by addition of 1-(3-dimethylaminopropyl)-3-ethylcarbodiimide hydrochloride (4.2 g, 22 mmol). Then the mixture was heated up to 80 °C and stirred for 24 h. After that, the solvent was removed under vacuum. The residue was separated by column chromatography on silica gel (methanol / DCM = 1:10) to give the product as a colorless oil (2.1 g, 44%). ^1^H NMR (CDCl_3_) δ 4.22 (s, 2H), 3.65 (s, 6H), 2.61 (t, 2H), 2.52 (td, 2H), 2.01 (t, 1H).

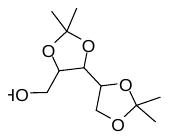

2,3:4,5-di-O-isopropylidene-D-xylitol (**4**)

This compound was synthesized similar to a procedure described in *Table 1* of a paper(*3*) with slight modification. To a suspension of xylitol (2.56 g, 16.8 mmol) in dry THF (48 mL) was added 2,2-dimethoxypropane (3.85 g, 37.0 mmol). After stirring the mixture at room temperature for 15 min, L-(-)-camphorsulfonic acid (390.3 mg, 1.68 mmol) was added. The stirring was continued for 24 h before the addition of K_2_CO_3_ (464.4 mg, 3.36 mmol). After removal of the solvent, the residue was dissolved in ethyl acetate, washed with water for three times. The organic phase was collected and the solvent was removed. The residue was separated and purified by column chromatography on silica gel (ethyl acetate / hexane =1:1) to obtain the product as a colorless oil (1.95 g, 51%). ^1^H NMR (CDCl_3_) δ 4.23-3.96 (m, 4H), 3.90-3.77 (m, 2H), 3.67-3.58 (m, 1H), 1.46- 1.40 (m, 9H), 1.37 (m, 3H).

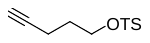

pent-4-ynyl tosylate (**5**)(*4*)

p-Toluenesulfonyl chloride (2.73 g, 14.3 mmol) was dissolved in dry DCM (35 mL) at 0 °C, 4-pentyn-1-ol (1 g, 11.9 mmol) and triethylamine (2.47 mL, 15.8 mmol) were then added dropwise. The mixture was stirred at room temperature for 20 h. After removal of the solvent, the residue was diluted with ether (100 mL) and filtered. The organic phase was collected and the solvent was removed. The residue was separated and purified by column chromatography on silica gel (ethyl acetate / hexane = 1:7) to give the product as a light yellow oil (1.37 g, 97%). ^1^H NMR (400 MHz, CDCl_3_) δ 7.82- -7.75 (d, 2H), 7.34 (d, 2H), 4.14 (t, 2H), 2.44 (s, 3H), 2.25 (td, 2H), 1.91-1.81 (m, 3H).

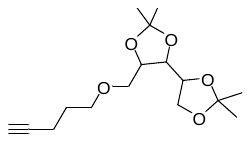

2,2,2',2'-tetramethyl-5-((pent-4-yn-1-yloxy)methyl)-4,4'-bi(1,3-dioxolane) (**6**)

At 0 °C, **4** (0.581 g, 2.5 mmol) was dissolved in dry THF (15 mL), and NaH (0.240 g, 10 mmol) was added under nitrogen. The mixture was stirred at room temperature for 1 h, and then **5** (1.192 g, 5 mmol) was gradually added. The mixture was further stirred at 50 °C for 36 h. Methanol (14 mL) and water (1 mL) were slowly added to quench the reaction, and the resulting mixture was stirred at 50 °C for 24 h for complete hydrolysis of the excess **5**. After removal of the volatile components, the residue was washed three times with water and ethyl acetate, and the organic phase was dried and filtered and then the solvent was removed. The residue was separated and purified by column chromatography on silica gel (ethyl acetate / hexane = 1:5) to give the product as a colorless oil (430 mg, 58%). ^1^H NMR (CDCl_3_) δ 4.17 (td, 1H), 4.04 (tt, 2H), 3.90 (dd, 1H), 3.84 (dd, 1H), 3.61-3.52 (m, 4H), 2.27 (td, 2H), 1.93 (t, 1H), 1.46-1.40 (m, 9H), 1.36 (m, 3H). ^13^C NMR (CDCl_3_) δ 109.9, 109.8, 83.9, 78.6, 76.6, 75.8, 71.7, 70.2, 68.7, 65.8, 28.5, 27.2, 27.1, 26.4, 25.6, 15.3. HRMS (ESI), m/z calcd. for C_16_H_26_O_5_ [M + Na]^+^: 321.1672; found 321.1679.

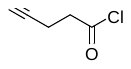

pent-4-ynoyl chloride (**7**)

4-Pentynoic acid (3.53 g, 36 mmol) and oxalyl chloride (5.48 g, 43.2 mmol) were dissolved in dry DCM (20 mL) and refluxed at 70 °C for 1 h. After removal of the solvent, a dark-red oily product (4.19 g, 100%) was obtained and directly used for the following esterification step.

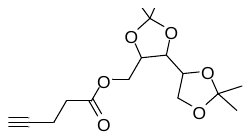

(2,2,2',2'-tetramethyl-[4,4'-bi(1,3-dioxolan)]-5-yl)methyl pent-4-ynoate (**8**)

**4** (1.4 g, 6 mmol) was added to a mixture solvents of N,N-diisopropylethylamine (4 mL) and dry DCM (20 mL). The solution was cooled down to 0 °C, and **7** (2.10 g, 18 mmol) was added dropwise. After the mixture was stirred at room temperature for 15 h, it was washed three times with saturated aqueous potassium carbonate solution, and the organic phase was collected, dried and filtered, and the solvent was removed. The residue was separated and purified by column chromatography on silica gel (ethyl acetate / hexane = 1:4) to give the product as a light yellow oil (461 mg, 25%). ^1^H NMR (CDCl_3_) δ 4.38-4.29 (m, 1H), 4.23-4.11 (m, 3H), 4.06 (dd, 1H), 3.93-3.84 (m, 2H), 2.64-2.58 (m, 2H), 2.54-2.48 (m, 2H), 1.97 (t, 1H), 1.46-1.35 (m, 12H). ^13^C NMR (151 MHz, CDCl_3_) δ 171.6, 110.3, 110.0, 82.4, 77.5, 69.8, 69.4, 69.3, 65.6, 64.8, 64.6, 64.1, 33.3, 27.2, 27.1, 27.1, 27.0, 26.3, 25.5, 18.5, 14.4. HRMS (ESI), m/z calcd. for C_16_H_24_O_6_ [M + Na]^+^: 335.1465; found 335.1472.

**General Procedure A** for the preparation of enediynes with (masked) hydroxyl group(s).

Under a nitrogen atmosphere, compound **2** (252 mg, 0.6 mmol), NHC-PdCl_2_-3-chloropyridine catalyst(*5*) (41 mg, 0.06 mmol), CuI (46 mg, 0.24 mmol), and diisopropylethylamine (0.3 mL, 1.8 mmol) were successively added into a mixture of dry THF (2 mL) and toluene (4 mL). The terminal alkyne(s) (0.9 mmol each or 1.8 mmol in the synthesis of **EDY-2** and **EDY-3**) in THF (0.6 mL) was then added dropwisely. The mixture was stirred at room temperature and monitored with TLC. After complete consumption of **2**, the mixture was purified through column chromatograph on silica gel to give the desired compound.

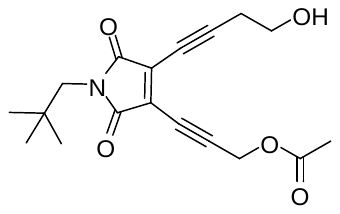
 **EDY-1**

3-(4-(4-hydroxybut-1-yn-1-yl)-1-neopentyl-2,5-dioxo-2,5-dihydro-1H-pyrrol-3-yl) prop-2-yn-1-yl acetate

Prepared according to the **General Procedure A**, isolated in 27% yield as a dark red oil. ^1^H NMR (CDCl_3_) δ 4.97 (s, 2H), 3.87 (t, J = 6.0 Hz, 2H), 3.34 (s, 2H), 2.85 (t, J = 6.0 Hz, 2H), 2.14 (s, 3H), 0.91 (s, 9H). ^13^C NMR (CDCl_3_) δ 170.3, 167.6, 130.4, 127.0, 110.5, 102.2, 76.2, 73.0, 60.4, 52.6, 50.3, 33.7, 28.0, 25.1, 20.8. HRMS (ESI), m/z calcd. for C_18_H_21_NO_5_Na [M + Na]^+^: 354.1317, found 354.1316.

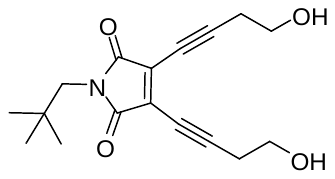
 **EDY-2**

3,4-bis(4-hydroxybut-1-yn-1-yl)-1-neopentyl-1H-pyrrole-2,5-dione

Prepared according to the **General Procedure A**, isolated in 68% yield as a yellow oil. ^1^H NMR (CDCl_3_) δ 3.85 (t, J = 6.0 Hz, 4H), 3.33 (s, 2H), 2.83 (t, J = 6.0 Hz, 4H), 0.90 (s, 9H). ^13^C NMR (CDCl_3_) δ 167.1, 127.5, 108.0, 72.0, 59.3, 49.12, 32.5, 26.9, 23.9. HRMS (ESI), m/z calcd. for C_17_H_21_NO_4_Na [M + Na]^+^: 326.1368, found 326.1366.

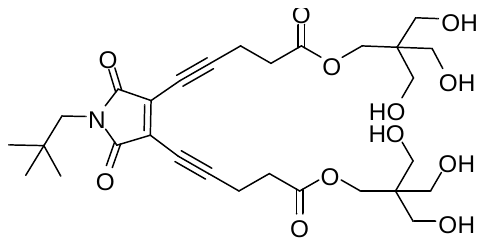
 **EDY-3**

bis(3-hydroxy-2,2-bis(hydroxymethyl)propyl)5,5'-(1-neopentyl-2,5-dioxo-2,5-dihydro-1H-pyrrole-3,4-diyl) bis(pent-4-ynoate)

Prepared according to the **General Procedure A**, isolated in 64% yield as a red oil. ^1^H NMR (CD_3_OD) δ 4.15 (s, 4H), 3.59 (s, 12H), 3.30 (s, 2H), 2.91 (t, J = 7.1 Hz, 4H), 2.73 (t, J = 7.1 Hz, 4H), 0.90 (s, 9H). ^13^C NMR (CD_3_OD) δ 171.8, 167.9, 128.1, 109.4, 71.3, 63.3, 61.0, 47.6, 44.9, 32.9, 32.4, 26.9, 15.5. HRMS (ESI), m/z calcd. for C_29_H_41_NO_12_Na [M + Na]^+^: 618.2526, found 618.2527.

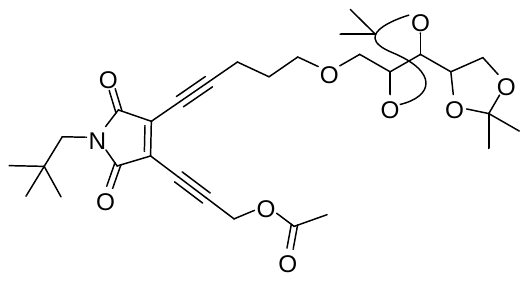
 **EDY-4**

3-(1-neopentyl-2,5-dioxo-4-(5-((2,2,2',2'-tetramethyl-[4,4'-bi(1,3-dioxolan)]-5-yl)methoxy)pent-1-yn-1-yl)-2,5-dihydro-1H-pyrrol-3-yl)prop-2-yn-1-yl acetate

Prepared according to the **General Procedure A**, isolated in 11% yield as a red oil. ^1^H NMR (CDCl_3_) δ 4.98 (s, 2H), 4.21-4.15 (m, 1H), 4.08 (dt, 1H), 4.04 (dd, 1H), 3.90 (dd, 1H), 3.87-3.82 (m, 1H), 3.63 (t, 2H), 3.59 (d, 2H), 3.34 (s, 2H), 2.69 (t, 2H), 2.13 (s, 3H), 1.93 (m, 2H), 1.45-1.37 (m, 12H), 0.91 (s, 9H). ^13^C NMR (CDCl_3_) δ 170.1, 167.8, 167.585, 130.5, 126.4, 109.9, 109.8, 78.5, 76.6, 76.4, 75.8, 72.1, 71.8, 69.9, 65.8, 52.6, 50.3, 33.7, 28.2, 28.0, 27.2, 27.1, 26.4, 25.6, 20. 8, 17.6. HRMS (ESI), m/z calcd. for C_30_H_41_NO_9_ [M + Na]^+^: 582.2674; found 582.2677.

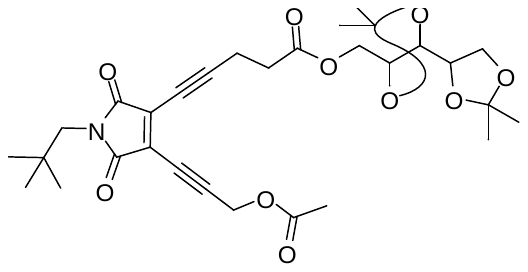
 **EDY-5**

((4S,5S)-2,2,2',2'-tetramethyl-[4,4'-bi(1,3-dioxolan)]-5-yl)methyl 5-(4-(3-acetoxyprop-1-yn-1-yl)-1-neopentyl-2,5-dioxo-2,5-dihydro-1H-pyrrol-3-yl)pent-4-ynoate

Prepared according to the **General Procedure A**, isolated in 25% yield as a red oil. ^1^H NMR (CDCl_3_) δ 4.98 (s, 2H), 4.35 (m, 1H), 4.23-4.10 (m, 2H), 3.92-3.84 (m, 2H), 3.33 (s, 2H), 2.92 (dt, 2H), 2.80-2.71 (m, 2H), 2.14 (s, 3H), 1.46-1.35 (m, 12H), 0.90 (s, 9H). ^13^C NMR (CDCl_3_) δ 171.1, 170.1, 167.6, 167.4, 130.0, 127.1, 110.4, 110.0, 77.5, 76.2, 75.4, 74.9, 72.2, 65.7, 64.9, 52.6, 50.4, 33.7, 32.7, 28.0, 27.2, 27.1, 26.3, 25.4, 20.8, 16.4. HRMS (ESI), m/z calcd. for C_30_H_39_NO_10_ [M + Na]^+^: 596.2466; found 596.2471.

**General Procedure B** for the preparation of hypersulfated enediynes.

The hypersulfation was conducted according to reported procedures(*6, 7*). A solution of **EDY-1** (1 equiv.) in anhydrous DMF (1 mL) maintained under an atmosphere of nitrogen was cooled to 0 °C and then treated dropwise with a solution of SO_3_·DMF (5 equiv. per OH) in anhydrous DMF (1 mL). The resulting mixture was stirred for 1 h and then allowed to warm to room temperature and stirred for an additional 2 h. Subsequently, the reaction flask was brought to -78 °C and quenched with sodium bicarbonate solution (10% w/v) until bubbling had ceased (pH = 7). After removal of the volatile components under reduced pressure (in a water bath maintained below 30 °C), the residue was washed with cold methanol, filtered and the filtrate was concentrated under room temperature. the methanol solution was then dropped into cold ether for re-precipitation, and the product was separated out by centrifugation.

**General Procedure C** for the removal of ketal mask groups in enediynes.

The **EDY-4** (50 mg, 0.09 mmol) was dissolved in DCM (1 mL), and then trifluoroacetic acid (0.2 mL) was slowly added. The mixture was stirred at room temperature and monitored by TLC for 2 h. After complete consumption of **EDY-4** and removal of the solvent, the product obtained in quantitative yield was directly used for the next step.

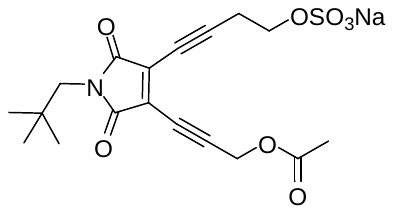
 **EDY-A**

Starting from **EDY-1**, prepared according to the **General Procedure B**, isolated in 99% yield as a dark-red oily solid. ^1^H NMR (D_2_O) δ 5.04 (s, 2H), 4.23 (t, J = 6.3 Hz, 2H), 3.34 (s, 2H), 3.03 (t, J = 6.2 Hz, 2H), 2.16 (s, 3H), 0.87 (s, 9H). ^13^C NMR (D_2_O) δ 178.6, 172.9, 169.0, 129.8, 127.1, 109.4, 102.7, 65.7, 52.8, 48.8, 32.8, 27.1, 21.5, 20.2. HRMS (ESI), m/z calcd. for C_18_H_20_NO_8_S^-^ [M]^-^: 410.0915, found 410.0912.

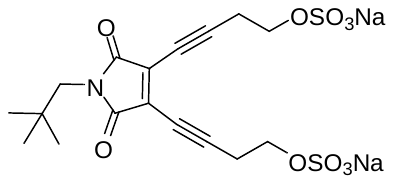
 **EDY-B**

Starting from **EDY-2**, prepared according to the **General Procedure B**, isolated in 68% yield as an orange oily solid. ^1^H NMR (D_2_O) δ 4.24 (t, J = 6.3 Hz, 4H), 3.32 (s, 2H), 3.03 (t, J = 6.3 Hz, 4H), 0.89 (s, 9H). ^13^C NMR (D_2_O) δ 169.8, 128.4, 108.2, 71.8, 65.9, 50.1, 32.7, 27.0, 20.9. HRMS (ESI), m/z calcd. for C_17_H_19_NNaO_10_S_2_^-^ [M]^-^: 484.0354, found 484.0354.

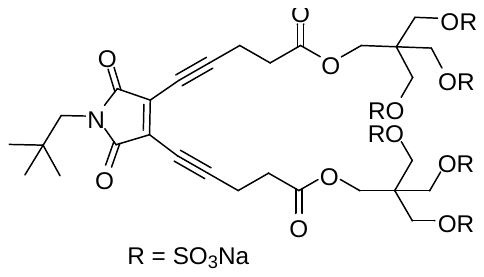
 **EDY-C**

Starting from **EDY-3**, prepared according to the **General Procedure B**. The solvent was removed under reduced pressure and the residue was then dialyzed in ultrapure water for 24 h (3 h, 9 h, 12 h, 23 h with water change) using a dialysis bag MD31 (MW100-500) to obtain a yellow transparent solution, which was then lyophilized to give a yellow solid in > 99% yield. ^1^H NMR (D_2_O) δ 4.18 (s, 4H), 4.04 (s, 12H), 3.22 (s, 2H), 2.91-2.85 (m, 4H), 2.74 (t, 4H), 0.83 (s, 9H). HRMS (ESI), m/z calcd. for C_29_H_35_NNa_4_O_30_S_6_^2-^ [M]^-^: 580.4585, found 580.4524.

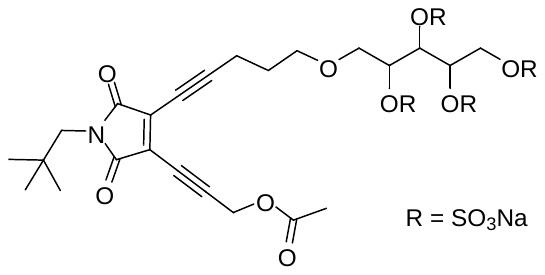
 **EDY-D**

Starting from **EDY-4**, hydrolyzed according to the **General Procedure C**, hypersulfated according to the **General Procedure B**. The solvent was removed by distillation under reduced pressure and the residue was then dialyzed in ultrapure water for 24 h (3 h, 9 h, 12 h, 23 h with water change) using a dialysis bag MD31 (MW100-500) to obtain a yellow transparent solution, which was then lyophilized to give a yellow solid in > 99% yield. ^1^H NMR (D_2_O) δ 5.10 (d, 2H), 5.02 (m, 1H), 4.97 (m, 1H), 4.49 (m, 1H), 4.40 (m, 1H), 3.95 (m, 1H), 3.77 (t, 2H), 3.36 (s, 2H), 2.76 (m, 2H), 2.20 (s, 3H), 1.96 (m, 2H), 0.92 (s, 9H).

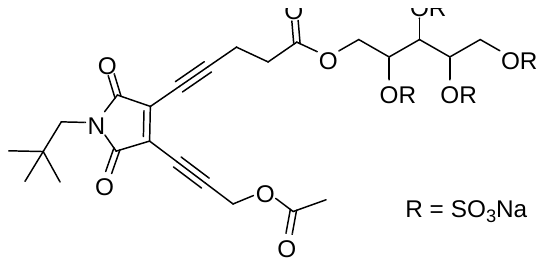
 **EDY-E**

Starting from **EDY-5**, hydrolyzed according to the **General Procedure C**, hypersulfated according to the **General Procedure B**. The solvent was removed by distillation under reduced pressure and the residue was then dialyzed in ultrapure water for 24 h (3 h, 9 h, 12 h, 23 h with water change) using a dialysis bag MD31 (MW100-500) to obtain a yellow transparent solution, which was then lyophilized to give a yellow solid in > 99% yield. ^1^H NMR (D_2_O) δ 5.00 (m, 5H), 4.53-4.28 (m, 4H), 3.07 (s, 2H), 2.82 (m, 2H), 2.19 (m, 5H), 0.92 (s, 9H).

1.4 **Radical generating property of enediyne antivirals**

A solution of N-tert-butyl-α-phenylnitrone (PBN, 100 mM) in DMSO was prepared in advance, then sulfated enediynes were added respectively to ensure their final concentration was 20 mM. The solution of PBN (100 mM) without addition of enediyne was used as the control. All samples were placed in a water bath at 37 °C and reacted for 12 h before the electron paramagnetic resonance (EPR) measurements. The EPR spectra were recorded with an X-band EMX-8/2.7C EPR spectrometer (Bruker, Germany).

2. **Materials and Methods for the biological tests for enediyne antivirals**
   2.1 **Cells and Viruses**

The Rhabdomyosarcoma (RD) cells line (ATCC: CCL-136), kidney of an African green monkey cells line VeroE6 (ATCC: CRL-1586) and human hepatocellular carcinoma cells line Huh-7 were cultured in Dulbecco’s modified Eagle’s medium (DMEM, Gibco, USA). Hela cells were obtained from the Chinese Academy of Science Cell Bank for Type Culture Collection (Shanghai, China). The human colorectal adenocarcinoma cells Caco-2(ATCC: HTB-37™) were transformed to overexpress hACE2 (hACE2/Caco-2). The cell line hACE2/Caco-2/Hela was cultured in DMEM too. All cells were cultured in medium supplemented with 10% fetal bovine serum (FBS), 100 U/ml penicillin, and 100 µg/ml streptomycin (Gibco, USA). Human coronavirus (HCoV)-OC43 (VR-1558) and HCoV-229E (VR-740) were kept by our laboratory (J.L.). HCoV-NL63 was gifted from Prof. Jincun Zhao (Guangzhou Medical University). SARS-CoV-2 Omicron variant (CCPM-B-V-049-2112-18) were provided by The Microorganisms and Viruses Culture Collection Center, Wuhan Institute of Virology, Chinese Academy of Sciences. The experiments involved SARS-CoV-2 were executed in biosafety level 3 (BSL-3) laboratory of Wuhan Institute of Virology, Chinese Academy of Sciences.

2.2 **The Viral S protein disintegration with enediyne antivirals**

The purified SARS-CoV-2 S protein (His-Avi) (0.1 mg/mL, CG202-01, Vazyme, China) was mixed with gradient concentration of enediyne (**EDY-A** ~ **EDY-E**) solutions. The mixtures were then incubated at 37 ℃ for 2 h (or 12 h). The degradation of protein was tested by SDS-PAGE and western blotting. The samples were separated by 10% SDS-PAGE and transferred to polyvinylidene difluoride (PVDF) membrane. The membranes were blocked with 5% skim milk at room temperature for 2 h and incubated with mouse antibody against His-tag(AE003, ABclonal, China) at 4 ℃ overnight. After washed by PBST, the membranes were incubated with horseradish peroxidase (HRP)-conjugated goat anti-mouse (A0216, Biyotime, China) or anti-rabbit antibodies (A0208, Biyotime, China) at room temperature for 1 h. Immunoblots were developed with the enhanced chemiluminescence reagents (180-5001, Tanon, China) and visualized by Tanon 4200SF imaging system.

2.3 **Cell Permeation of Enediynes**

The cellular permeation of enediynes in HeLa (or Vero E6) cells were examined with Confocal Laser Scanning Microscopy (CLSM). Hela cells (for example) were seeded in glass bottom confocal dishes at a density of 2 × 10^5^ per well in 2 mL of DMEM and incubated overnight. After removal of culture medium, cells were then incubated with hypersulfated or un-sulfated enediyne (1 μM) in 2 mL of DMEM. After 16 h of incubation at 37 ºC, the cells were washed three times with PBS. Subsequently, the cells were fixed with 2.5% glutaraldehyde at room temperature for 10 min, and permeabilized with 0.5% Triton X-100 for another 10 min. After that, the cells were washed with PBS, then 400 μL of propidium iodide (PI) solution (15 μg/mL) was added and the cells were cultured at 37 °C for 10 min, followed by washing with PBS for three times, and finally visualized by CLSM.

2.4 **Cytotoxicity of Enediynes**

Cytotoxicity of enediyne was tested based on CCK-8 cell counting kit (A311, Vazyme, China). The RD, Huh-7 and hACE2/Caco-2 cells were seeded separately into the 96-well microplates (10^4^ cells per well at 100 μL) and cultured at 37 ℃ overnight. Then added 14.3 μL 8-times working concentrations of enediyne solution to each well. The control group (adding 14.3 μL PBS) and blank wells were conducted at the same time. Each group is provided with three repeating wells. After incubated at 37 ℃ for 24 h, the supernatant was removed and replaced with 100 μL DMEM. After 10 μL CCK-8 solution was added to each well, the cells were incubated at 37 ℃ for 1 h. The absorbance at wavelength of 450 nm was detected with microplate reader (Biotek Synergy H1, USA). The cells viability rate was calculated as (OD value of experimental group -OD value of blank)/ (OD value of control group -OD value of blank) ×100%.

2.5 **Antiviral Activity of Sulfated Enediynes to Seasonal Coronavirus**

Seasonal coronavirus (HCoV-229E, HCoV-OC43 and HCoV-NL63) viral stock were diluted separately 50 times by PBS. The enediynes were serial diluted to different concentrations, mixed with HCoV and incubated at 37 ℃ for 2 h. To determine the viral titers, the mixtures were diluted from 10^-1^ to 10^-6^ and added to the monolayer Huh-7, RD and Caco-2 cells, respectively. The control group of viruses without the enediyne was also set. After an incubation at 32 ℃ for 2 h, the supernatant was removed and culture medium (DMEM supplement with 2% heat-inactivated FBS, 100 U/mL penicillin, 100 μg/mL streptomycin and 1.2% Avicel) was layered on the cell. After 72 h culture, the layer was removed and the cells were fixed with 4% polyformaldehyde and stained with 1% crystal violet. After staining at room temperature for 20 min, the plates were washed with running water to remove the dying liquid. The plaques were counted after the plates were dried at room temperature. The inhibition rate of enediyne against seasonal coronavirus was calculated as (number of plaques in control group - number of plaques in experimental group)/number of plaques in control group×100%

2.6 **Antiviral Activity of Sulfated Enediynes to SARS-CoV-2 Omicron Variant**

To determine the antiviral effects of the enediyne solutions, Vero E6 cells were seeded in 24 well culture plate at a density of 1×10^5^ cells per well and incubated for 16 hours before the antiviral assay was conducted. The enediyne solutions were serially diluted with PBS and then SARS-CoV-2 Omicron of 87,500 PFU in a volume of 175 μL were added. The mixtures were incubated at 37℃ for 2 h. Meanwhile, viruses were mixed with PBS and incubated at 4℃ for 2 h as the initial control. Then the mixtures were diluted by 10-fold, and the serially diluted virus stock were added into a 24-well culture plate containing Vero E6 cells. After infection at 37 ℃ for 2 h, the supernatant was discarded, and the cells were overlaid with medium containing 1% methylcellulose and incubated for the indicated time. Five days later, virus titers were determined by performing a plaque assay.

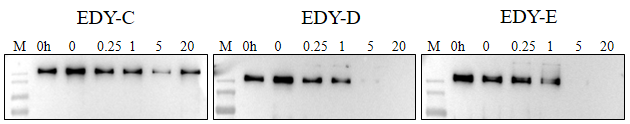

Figure S1. Western-Bolt analysis results of SARS-CoV-2 S protein after incubated with **EDY-C**, **EDY-D**, and **EDY-E** at 37 ℃ for 12 h, respectively. The lines marked with “0 h” mean the standard S protein samples.

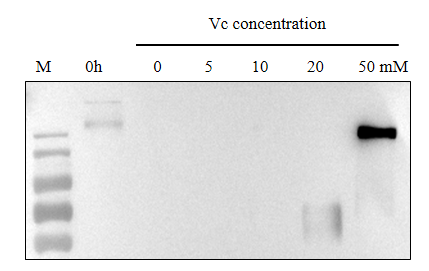

Figure S2. Western-Bolt analysis of SARS-CoV-2 S protein after incubated with **EDY-D** (5 mM) and series dilution of vitamin C. The line marked with “0 h” mean the standard S protein samples.

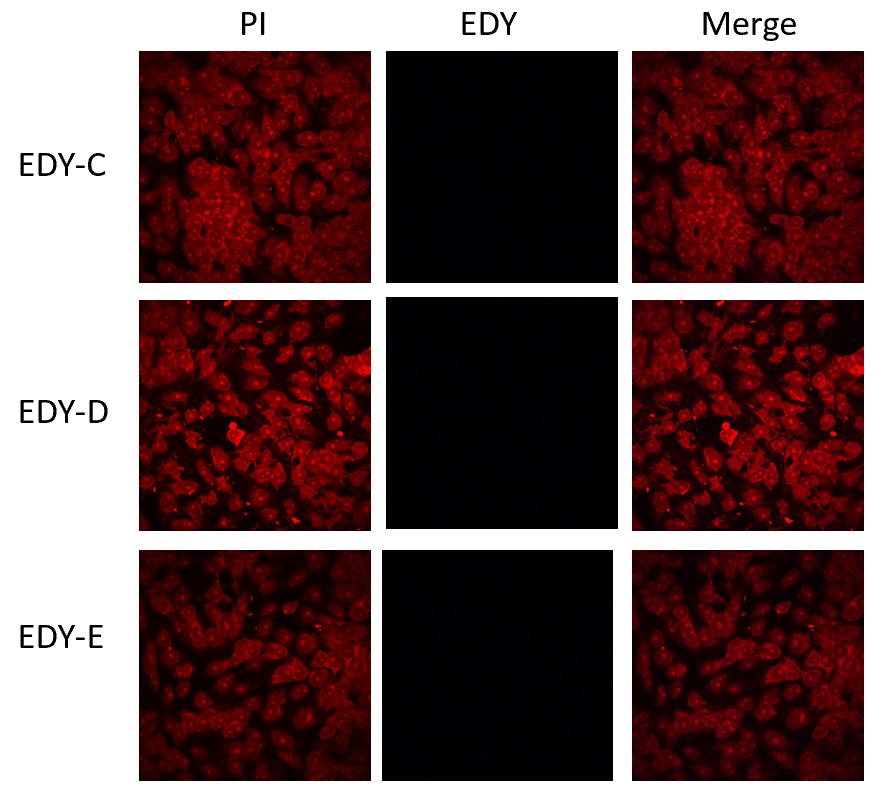

Figure S3. Confocal laser scanning microscopy images of Vero E6 cells treated with **EDY-C**, **EDY-D**, and **EDY-E**. Vero E6 cells are stained with propidium iodide (PI) showing red fluorescence while maleimide-based enediynes show intrinsic blue fluorescence (if there is any).

Table S1. Antiviral performance of enediynes against human coronaviruses

|  |  | EDY-A | EDY-B | EDY-C | EDY-D | EDY-E |
| --- | --- | --- | --- | --- | --- | --- |
| hCoV-229E  @Huh7 | CC50 (mM) | 0.76 | 0.85 | > 1.00 | 0.93 | 0.10 |
|  | EC50 (μM) | 13.50 | 87.75 | 38.48 | 3.86 | 14.78 |
|  | SI | 56 | 10 | > 26 | 242 | 7 |
| hCoV-NL63  @Caco2 | CC50 (mM) | 0.86 | 0.78 | > 1.00 | 9.92 | 0.11 |
|  | EC50 (μM) | 11.52 | 137.20 | 44.77 | 16.86 | 15.31 |
|  | SI | 75 | 6 | > 22 | 588 | 7 |
| hCoV-OC43  @RD | CC50 (mM) | 0.78 | 0.89 | > 1.00 | 0.74 | 0.08 |
|  | EC50 (μM) | 9.68 | 94.22 | 22.22 | 9.46 | 4.33 |
|  | SI | 80 | 9 | > 45 | 78 | 18 |
| SARS-CoV-2  @Vero E6 | CC50 (mM) | 0.78 | 0.70 | > 1.00 | 1.20 | 0.82 |
|  | EC50 (nM) | 99.60 | 796.40 | 256.20 | 56.19 | 132.20 |
|  | SI | 7831 | 878 | > 3903 | 21356 | 6209 |

NMR Characterization of enediynes and key precursors.

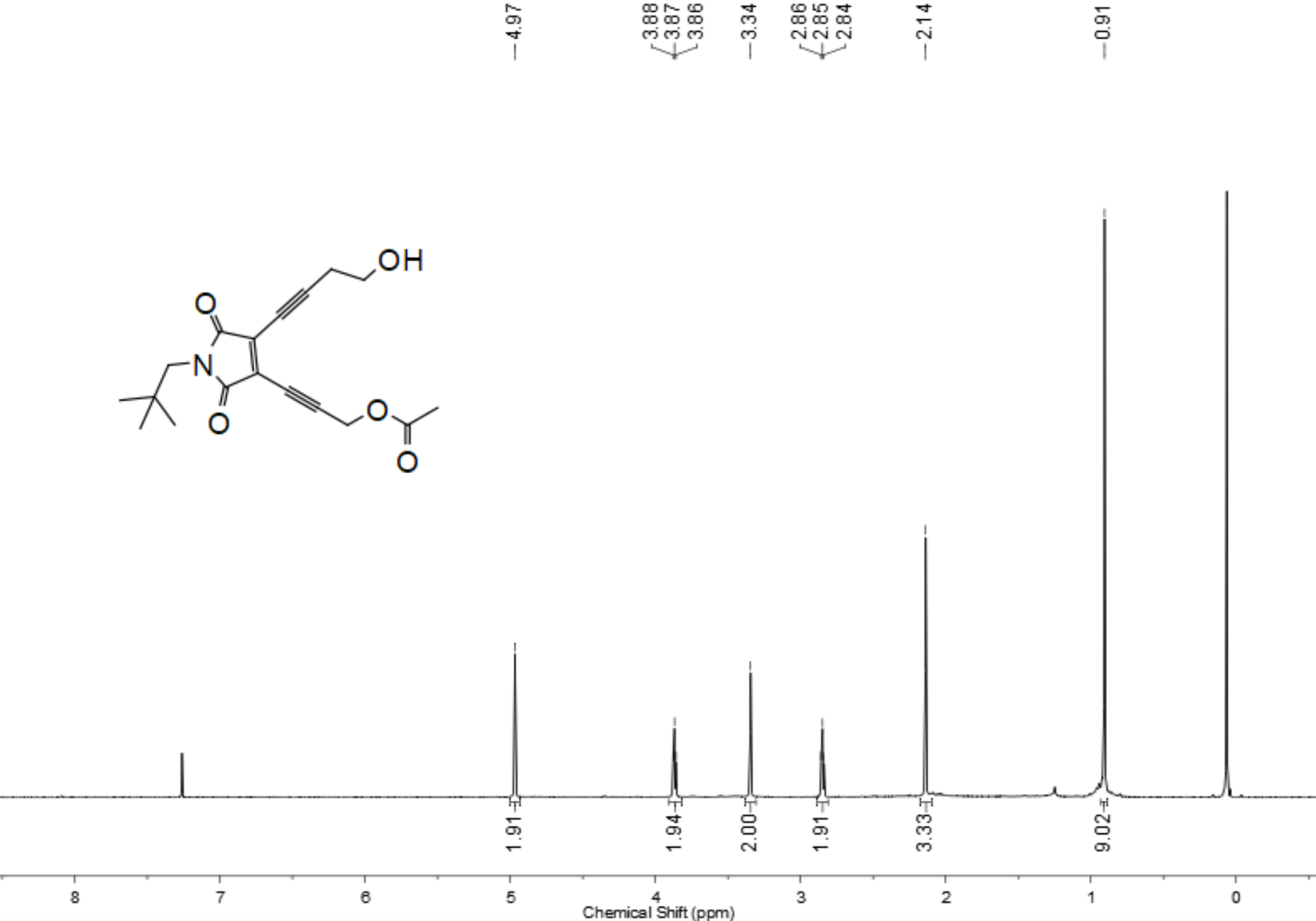

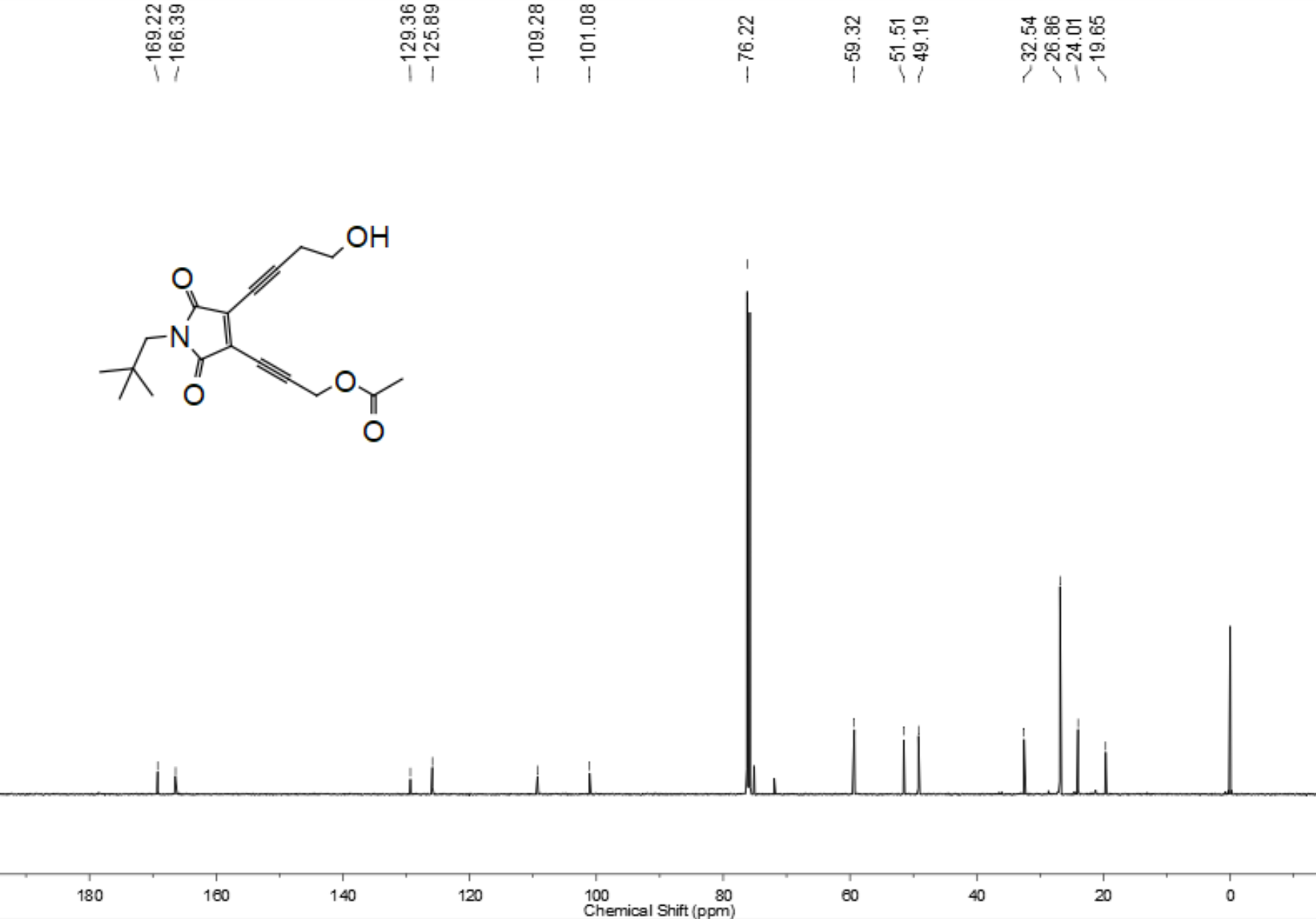

Figure S4 ^1^H NMR and ^13^C NMR spectra of **EDY-1**

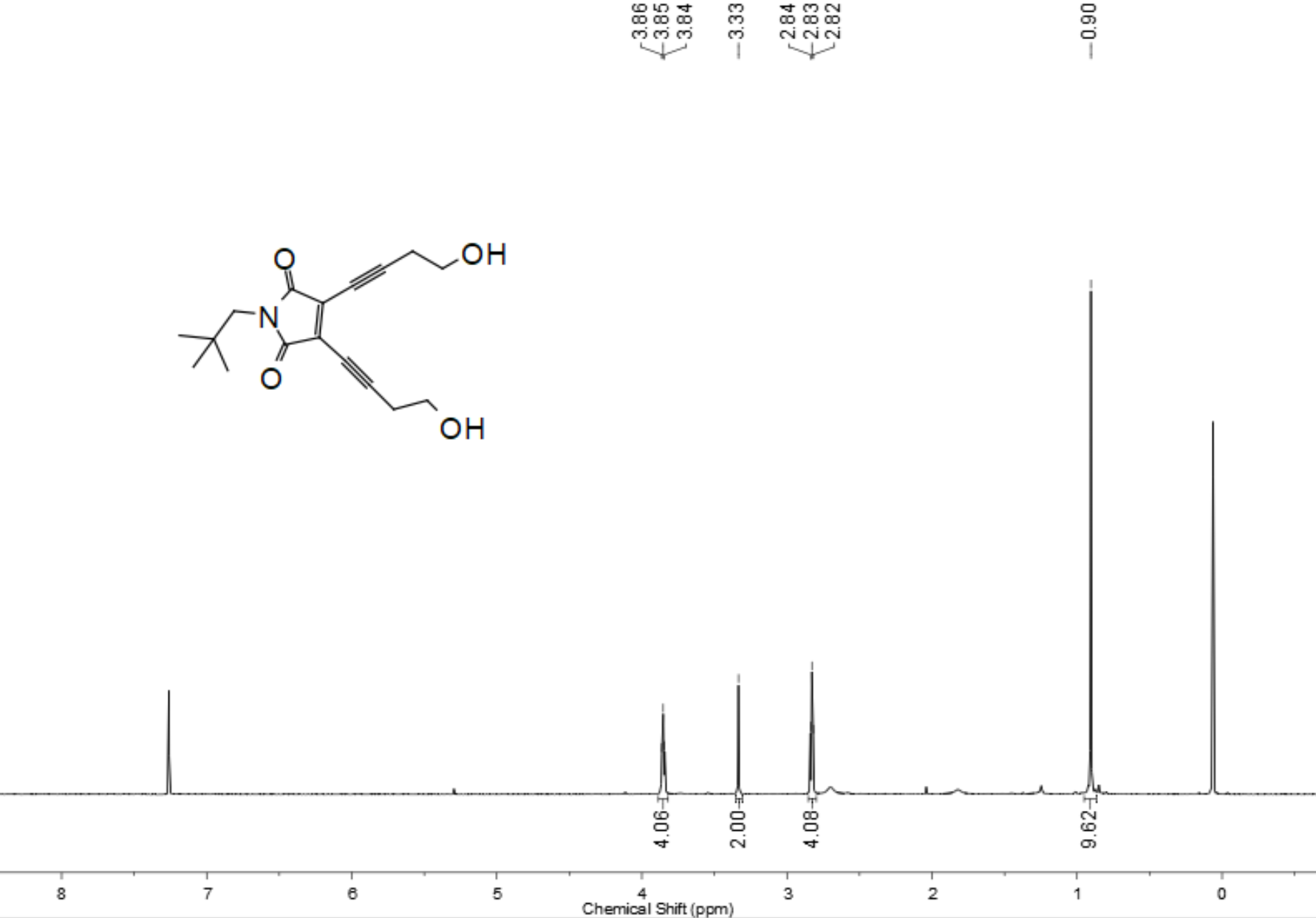

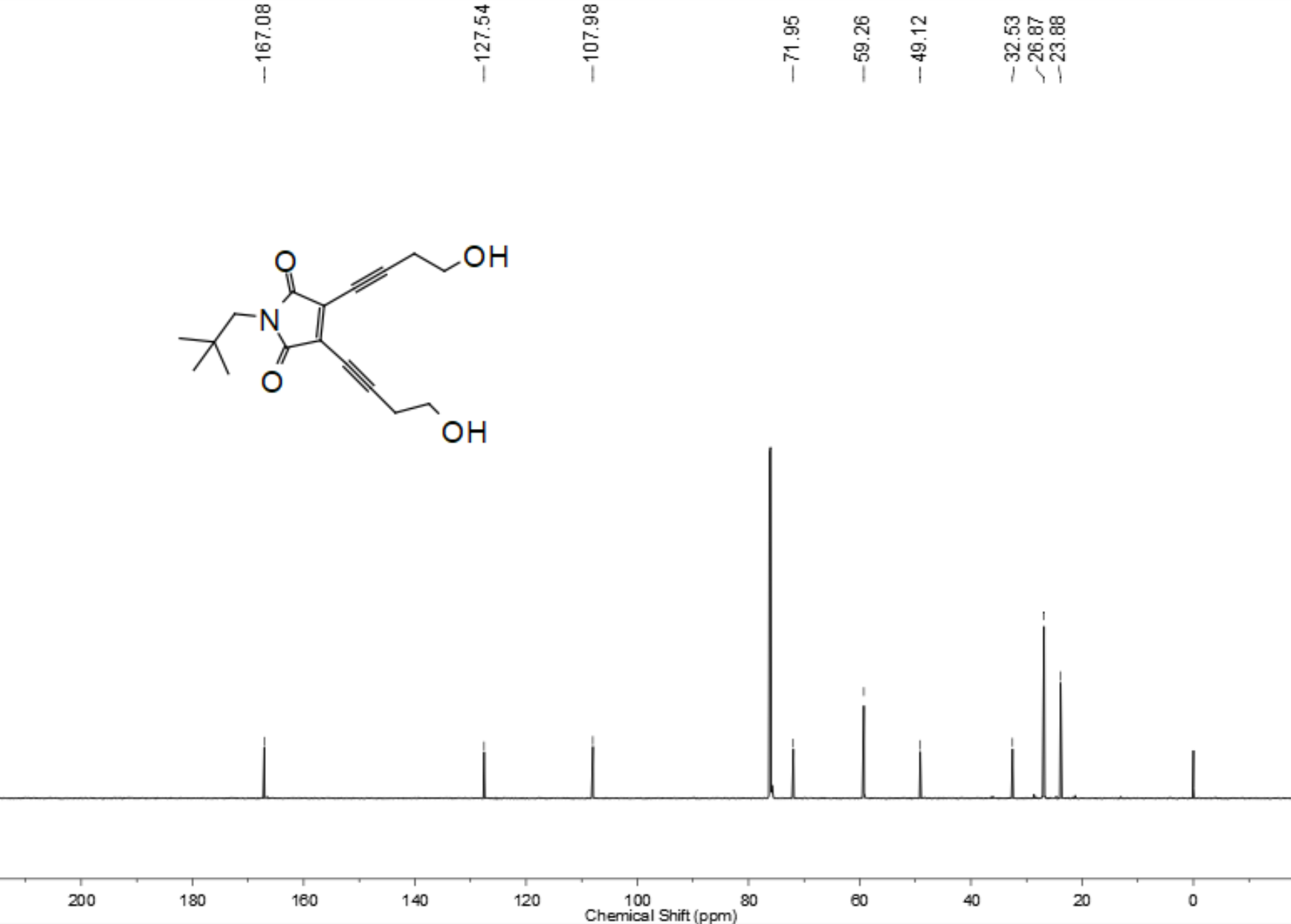

Figure S5 ^1^H NMR and ^13^C NMR spectra of **EDY-2**

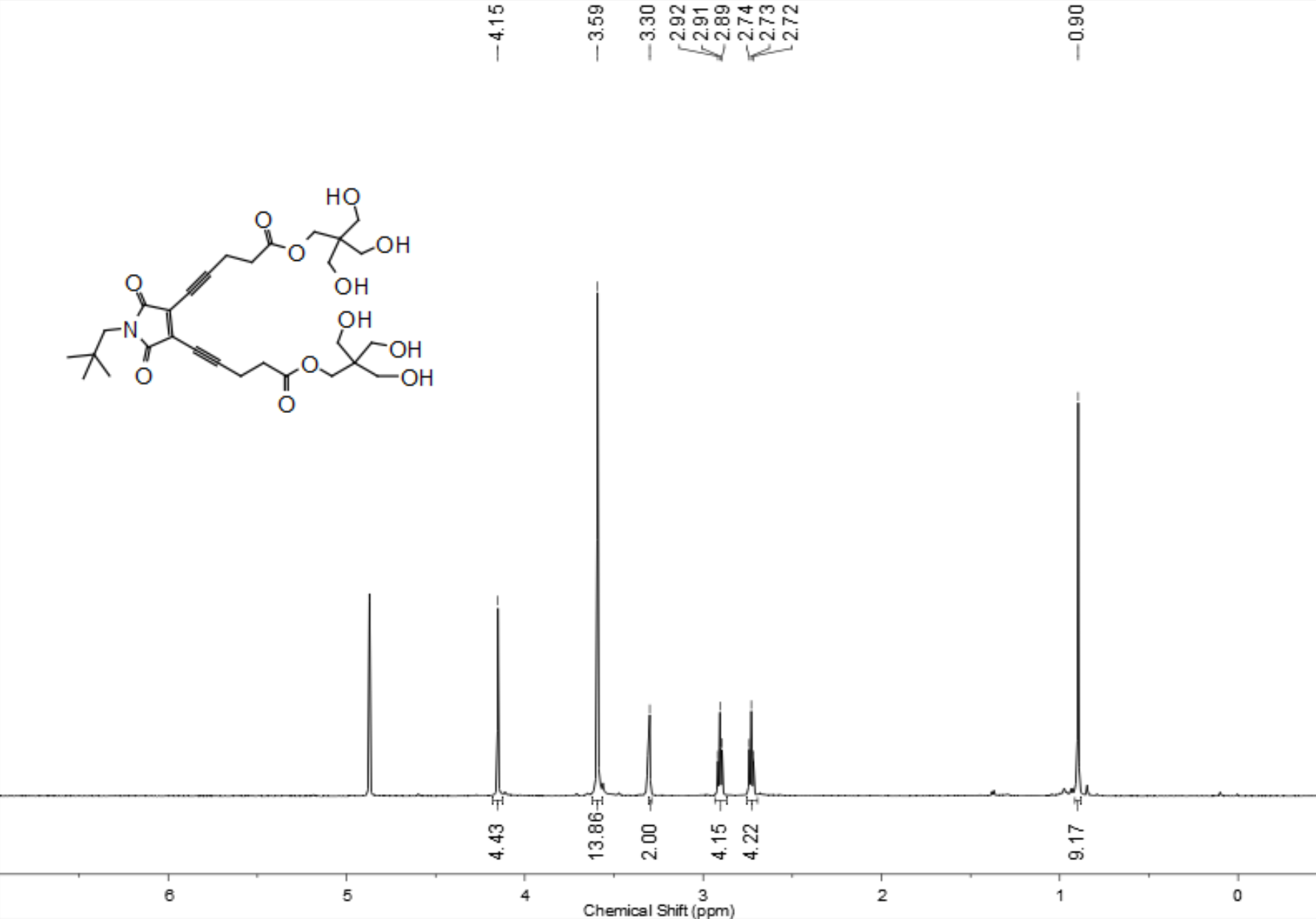

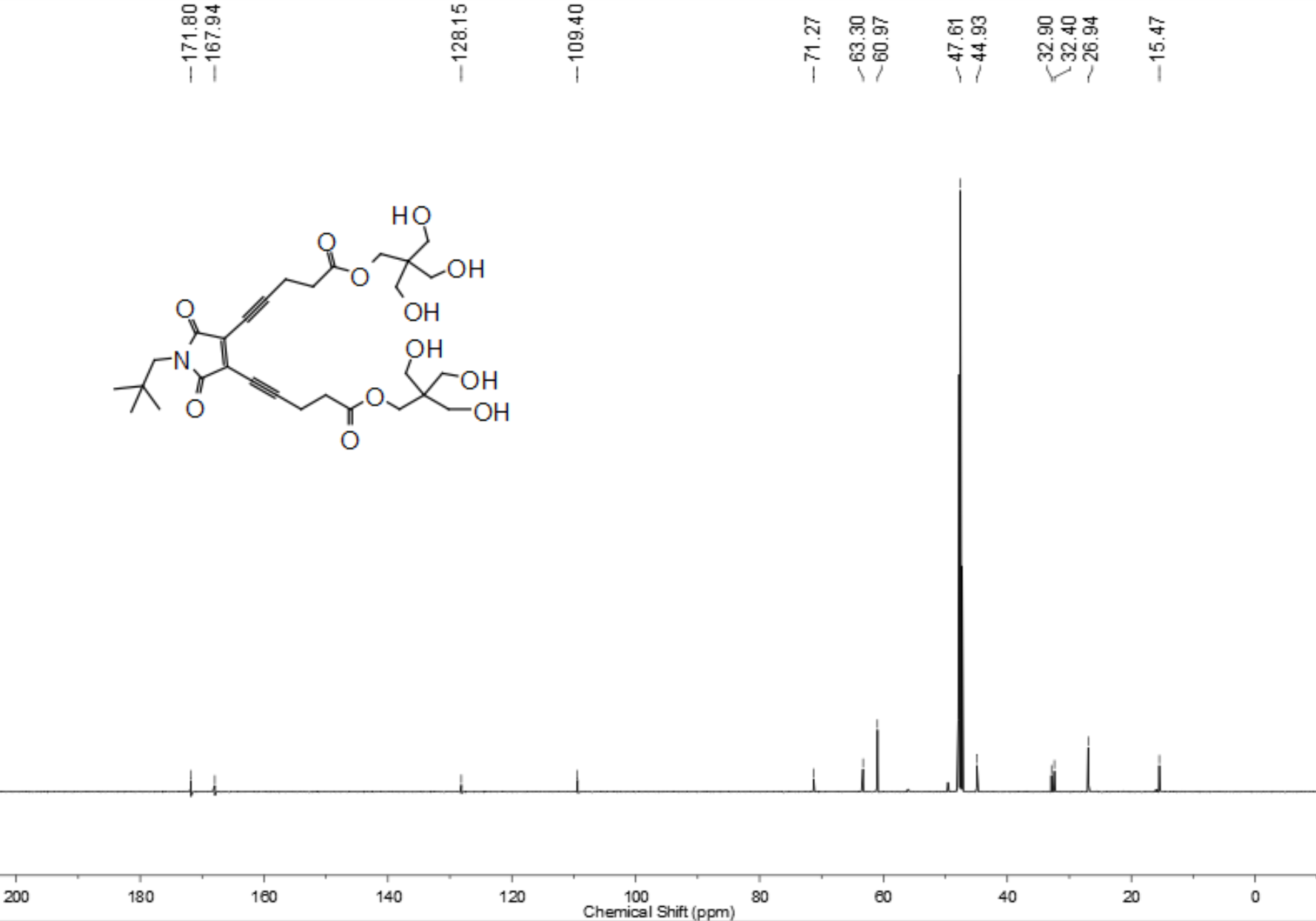

Figure S6 ^1^H NMR and ^13^C NMR spectra of **EDY-3**

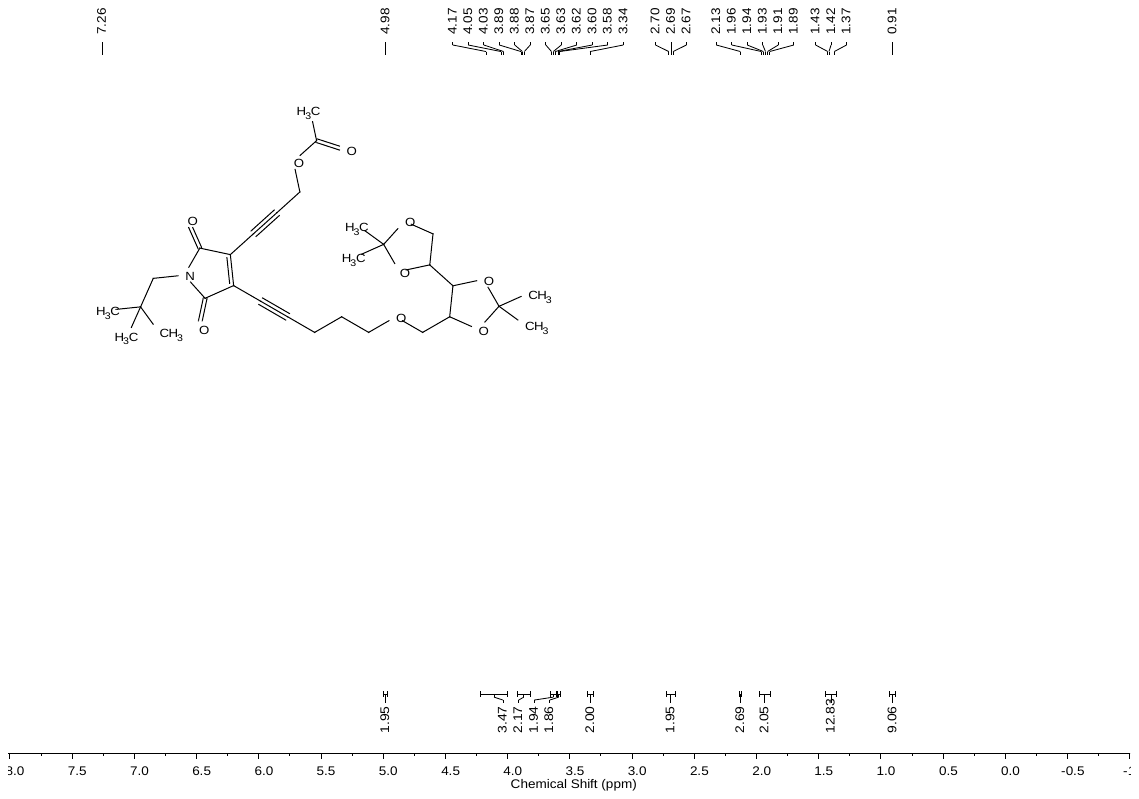

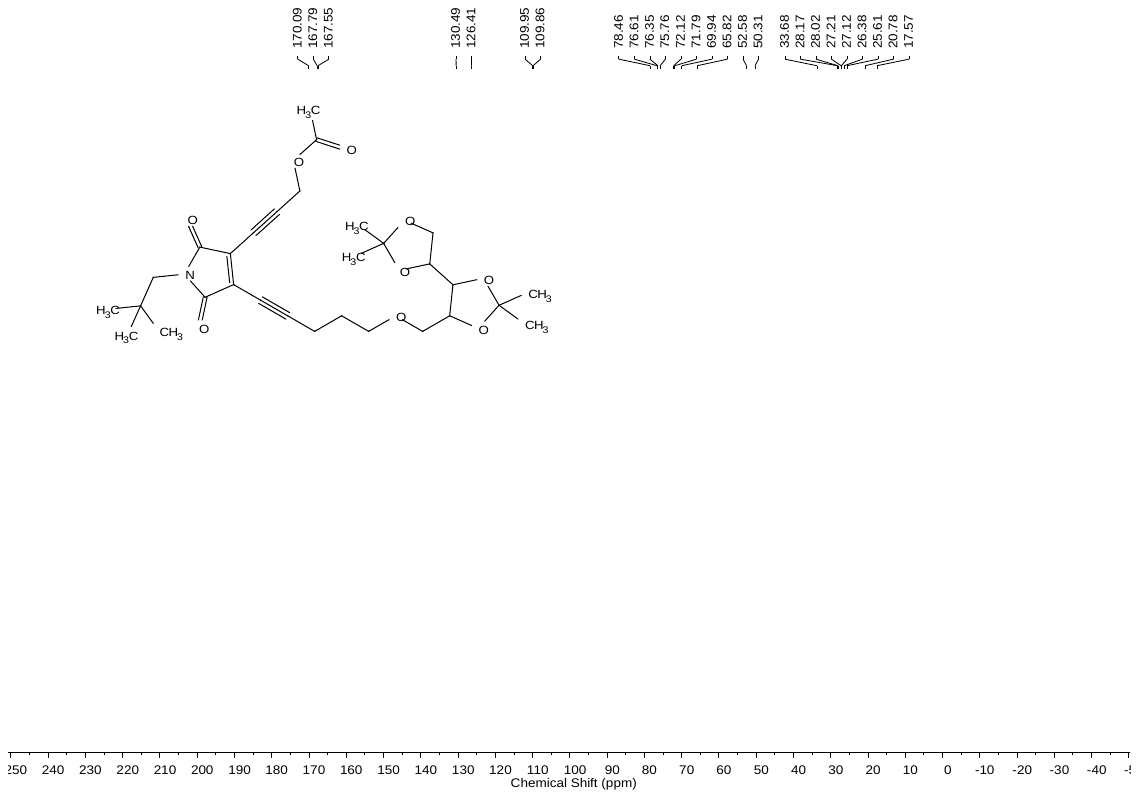

Figure S7 ^1^H NMR and ^13^C NMR spectra of **EDY-4**

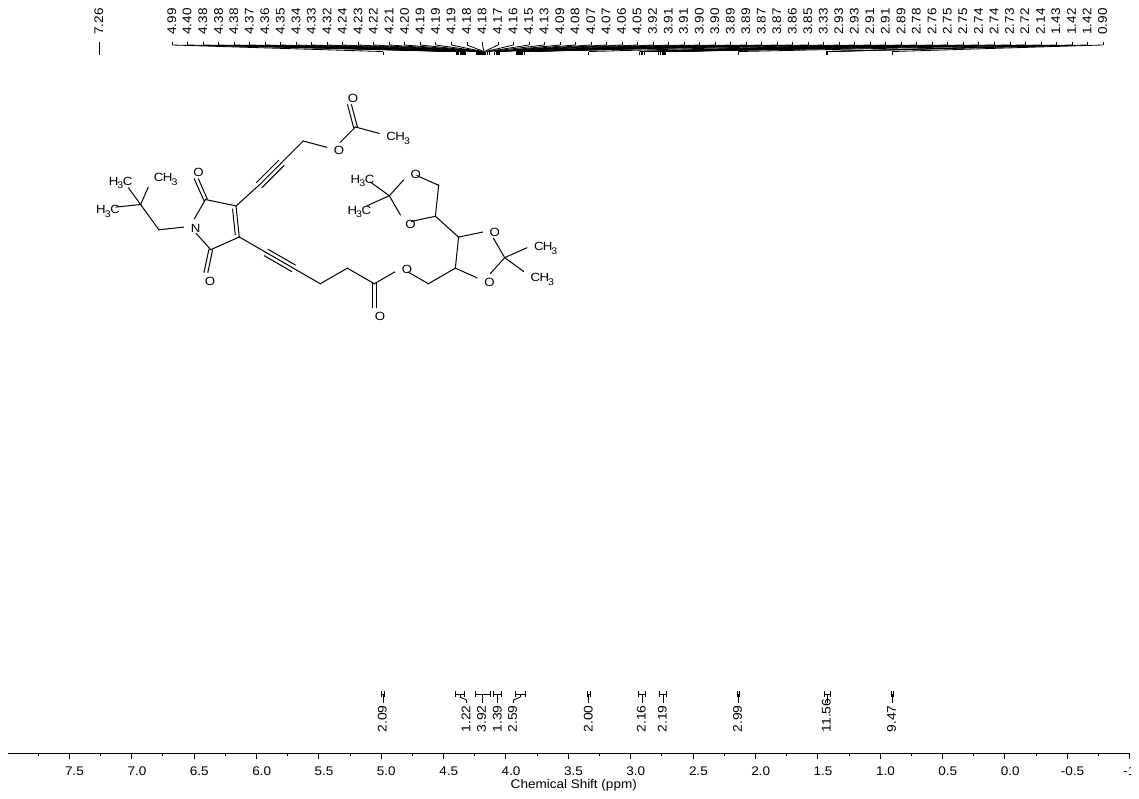

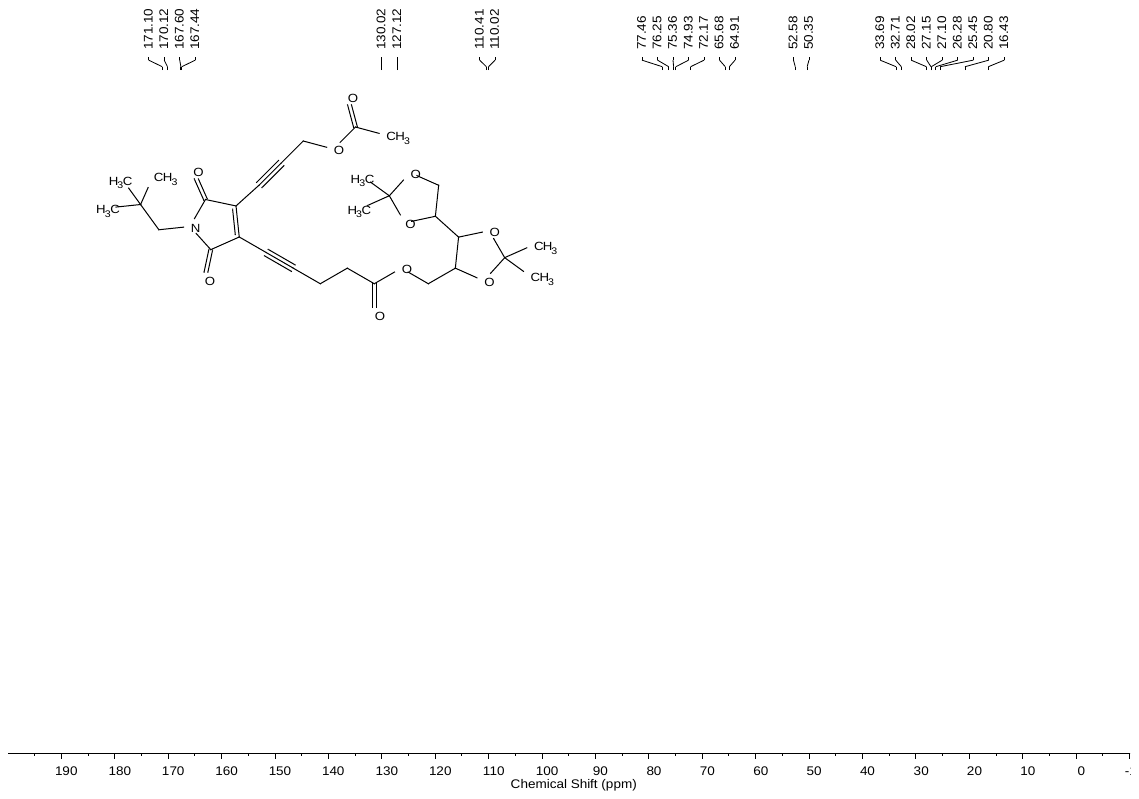

Figure S8 ^1^H NMR and ^13^C NMR spectra of **EDY-5**

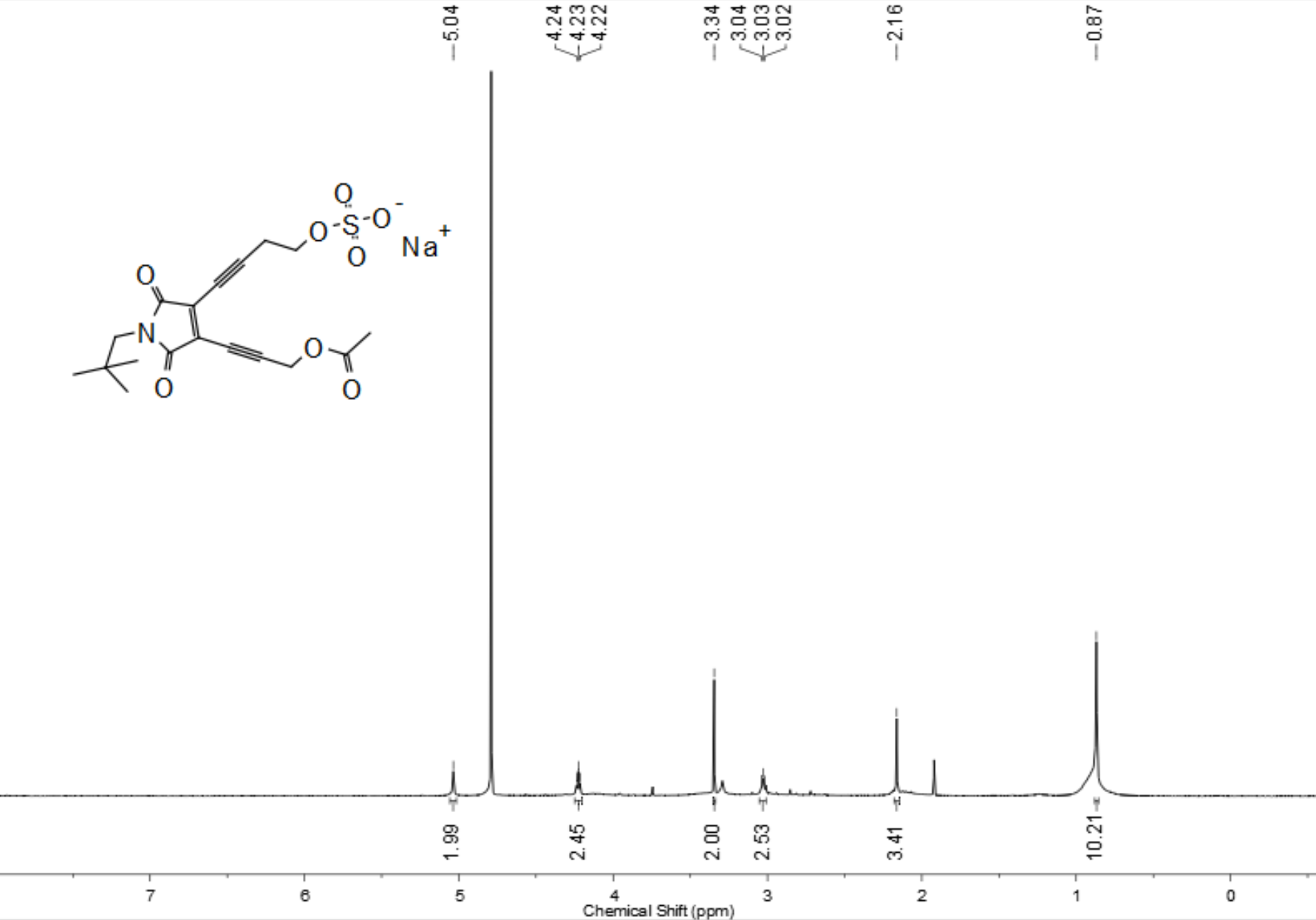

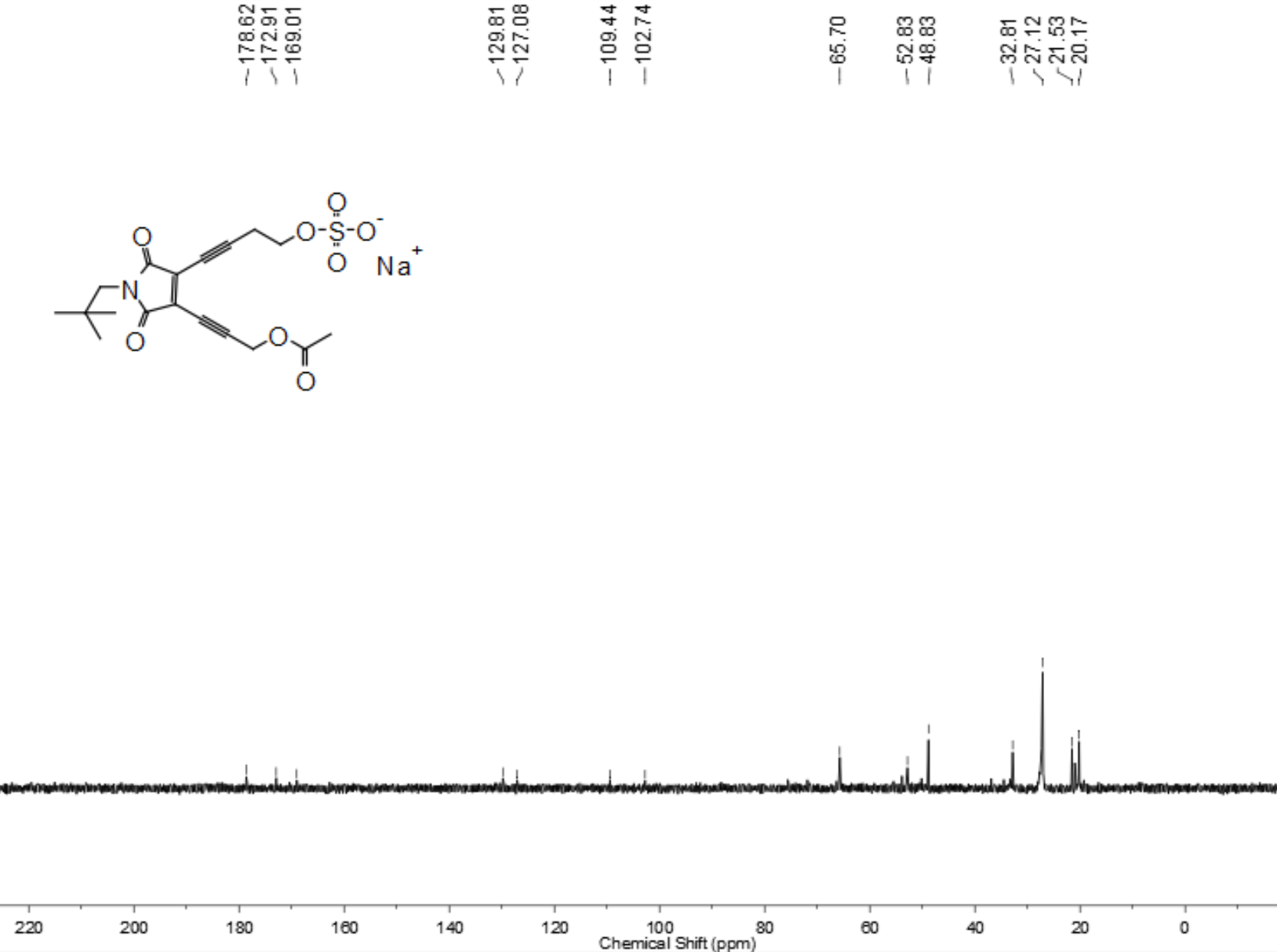

Figure S9 ^1^H NMR and ^13^C NMR spectra of **EDY-A**

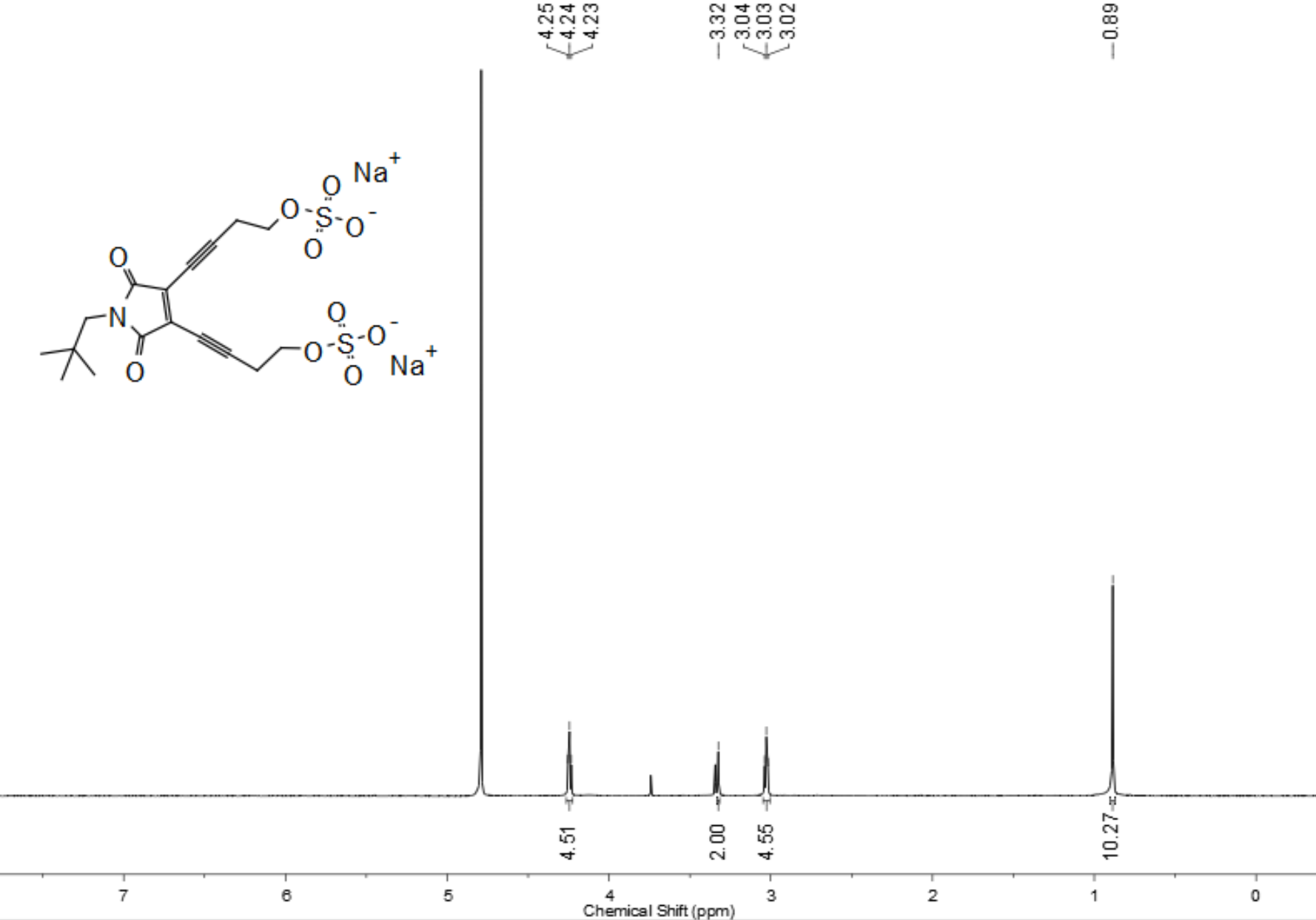

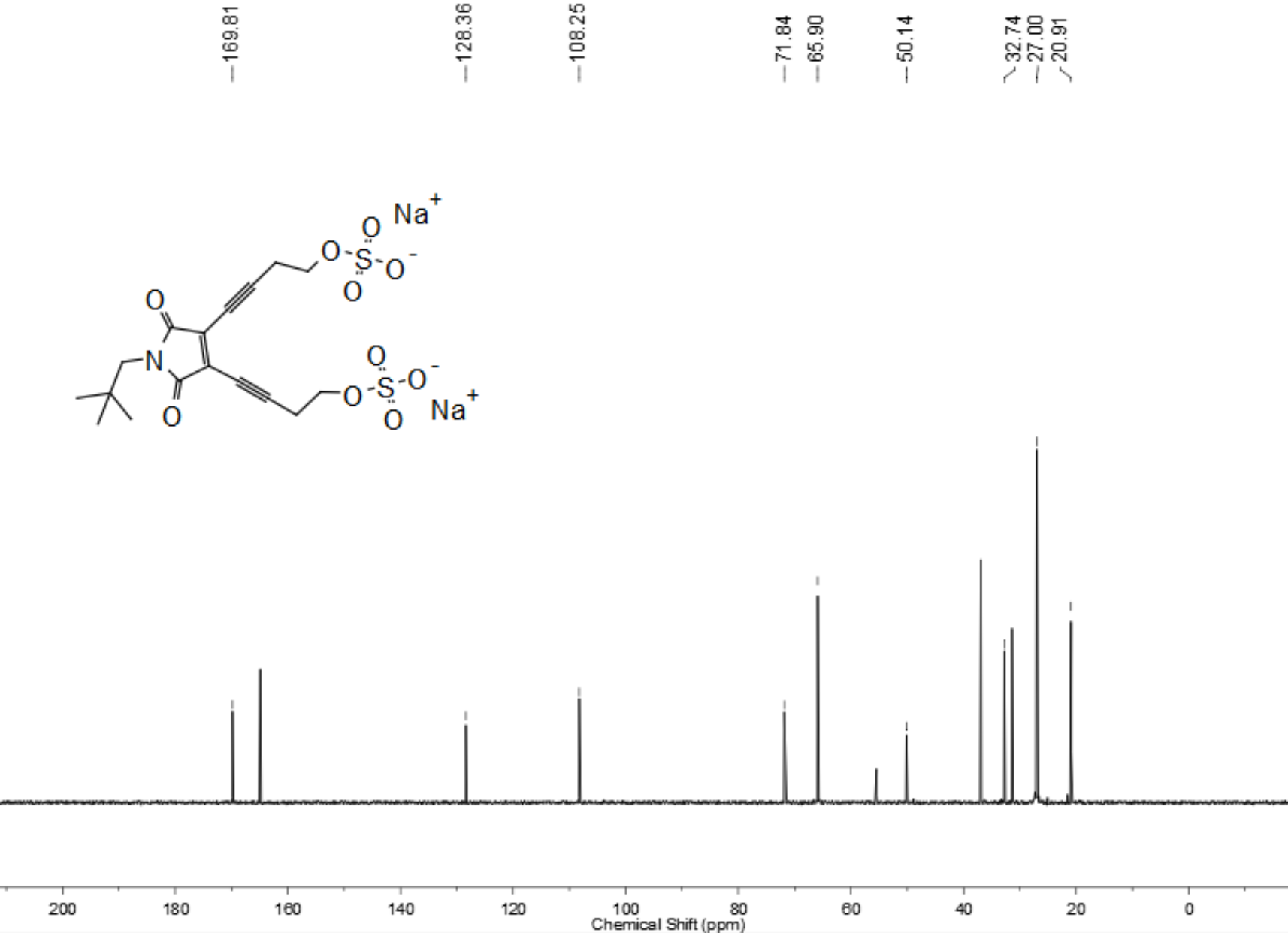

Figure S10 ^1^H NMR and ^13^C NMR spectra of **EDY-B**

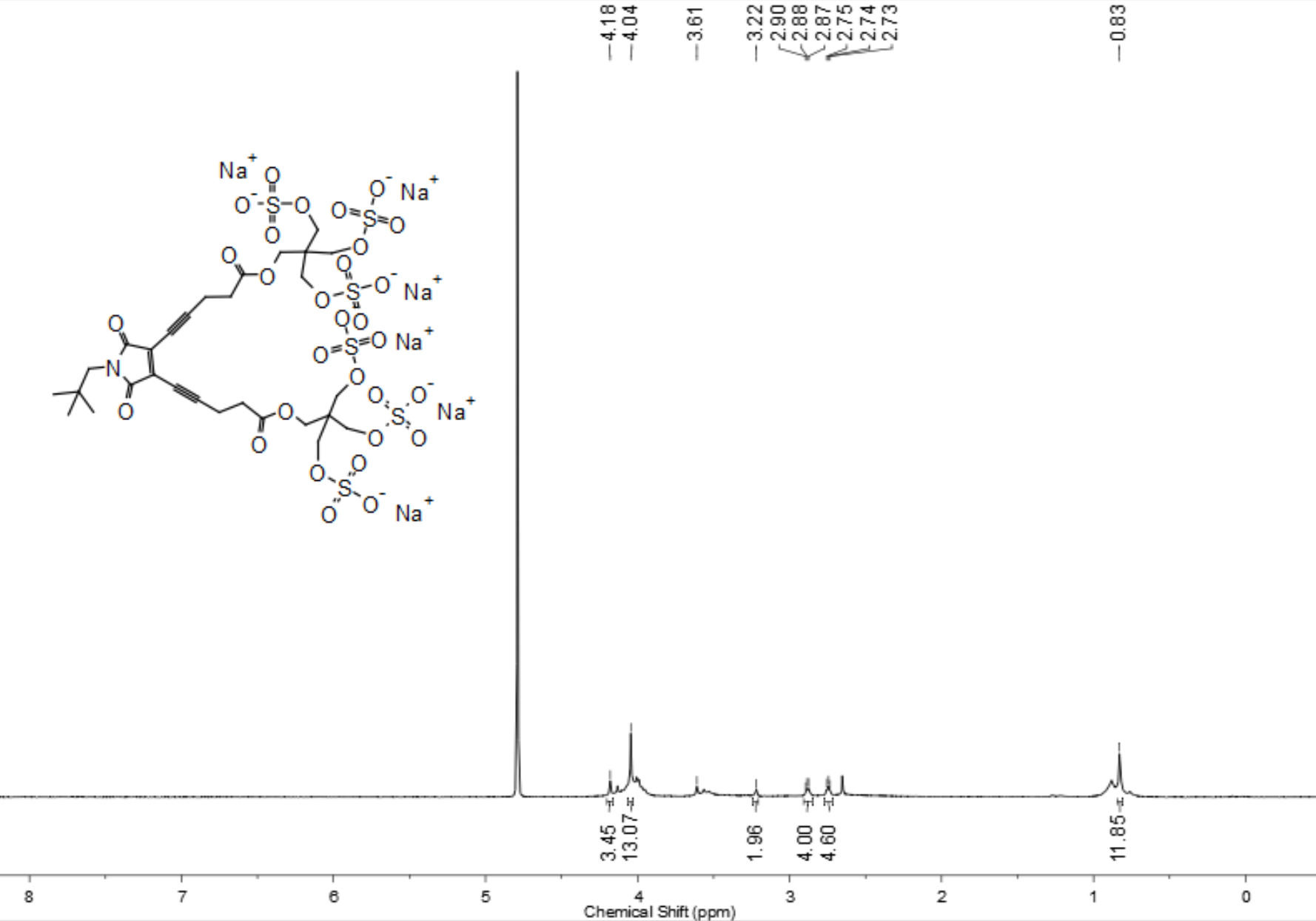

Figure S11 ^1^H NMR spectrum of **EDY-C**

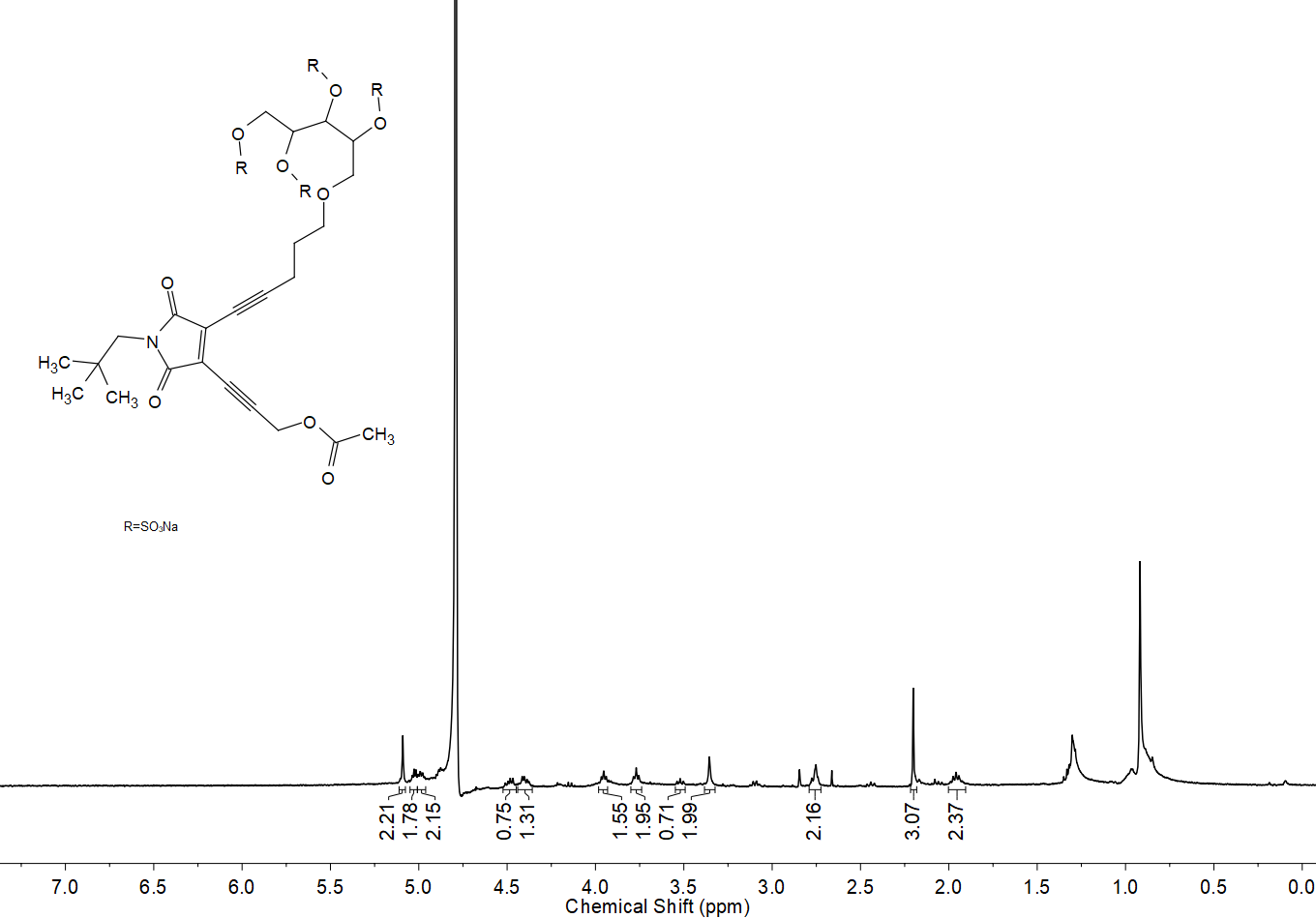

Figure S12 ^1^H NMR spectrum of **EDY-D**

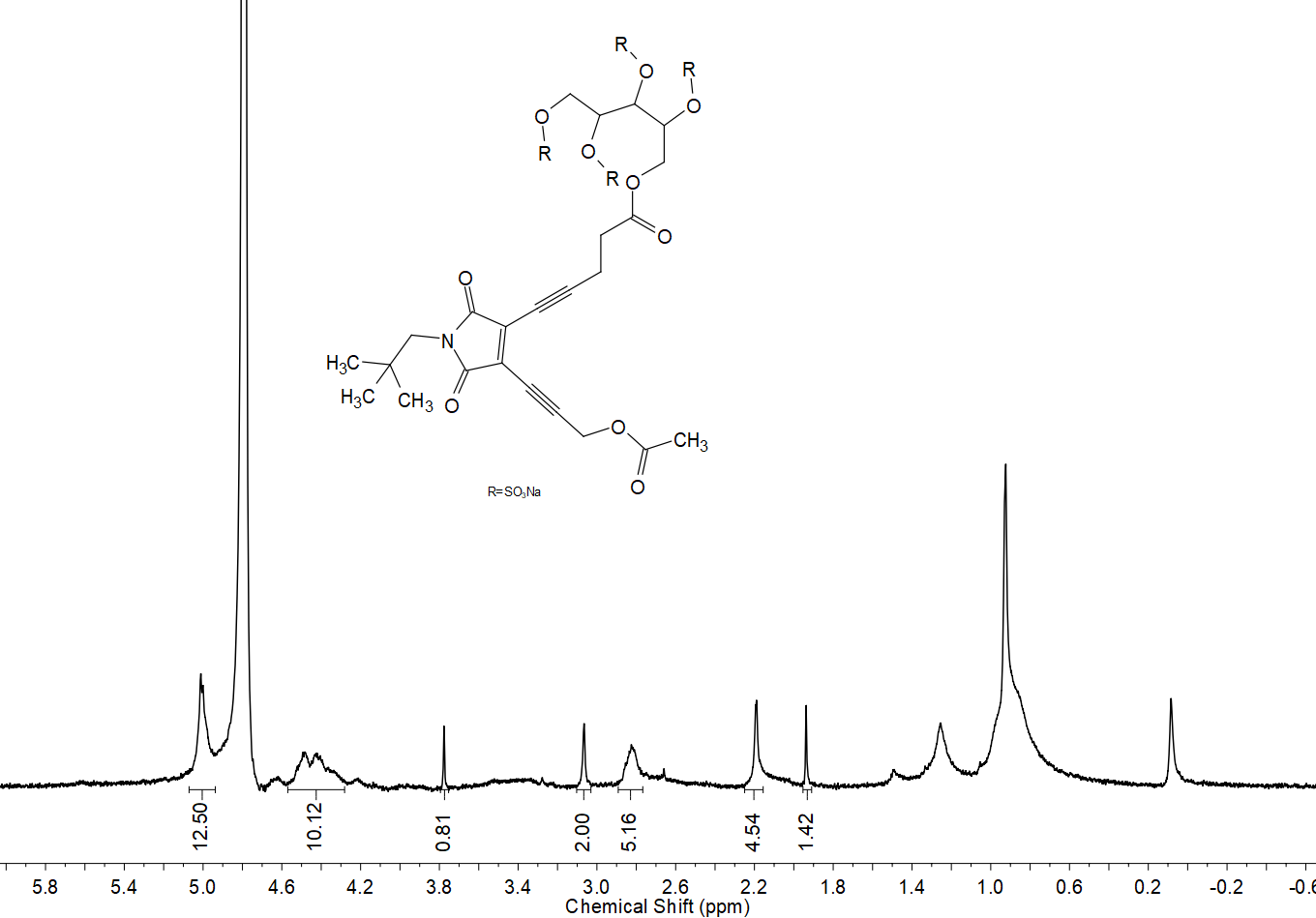

Figure S13 ^1^H NMR spectrum of **EDY-E**
